# Targeting fibrosis in the Duchenne Muscular Dystrophy mice model: an uphill battle

**DOI:** 10.1101/2021.01.20.427485

**Authors:** Marine Theret, Marcela Low, Lucas Rempel, Fang Fang Li, Lin Wei Tung, Osvaldo Contreras, Chih-Kai Chang, Andrew Wu, Hesham Soliman, Fabio M.V. Rossi

**Affiliations:** School of Biomedical Engineering and the Biomedical Research Centre, Department of Medical Genetics, 2222 Health Sciences Mall, Vancouver, BC, V6T 1Z3, Canada; Department of Pharmacology and Toxicology, Faculty of Pharmaceutical Sciences, Minia University, Minia, Egypt; Developmental and Stem Cell Biology Division, Victor Chang Cardiac Research Institute, Darlinghurst, NSW, 2010, Australia; Departamento de Biología Celular y Molecular and Center for Aging and Regeneration (CARE-ChileUC), Facultad de Ciencias Biológicas, Pontificia Universidad Católica de Chile, 8331150 Santiago, Chile

**Keywords:** drug screening, fibro/adipogenic progenitors, fibrosis, repair, skeletal muscle

## Abstract

**Aim:** Fibrosis is the most common complication from chronic diseases, and yet no therapy capable of mitigating its effects is available. Our goal is to unveil specific signallings regulating the fibrogenic process and to identify potential small molecule candidates that block fibrogenic differentiation of fibro/adipogenic progenitors.

**Method:** We performed a large-scale drug screen using muscle-resident fibro/adipogenic progenitors from a mouse model expressing EGFP under the *Collagen1a1* promotor. We first confirmed that the EGFP was expressed in response to TGFβ1 stimulation *in vitro*. Then we treated cells with TGFβ1 alone or with drugs from two libraries of known compounds. The drugs ability to block the fibrogenic differentiation was quantified by imaging and flow cytometry. From a two-rounds screening, positive hits were tested *in vivo* in the mice model for the Duchenne muscular dystrophy (mdx mice). The histopathology of the muscles was assessed with picrosirius red (fibrosis) and laminin staining (myofiber size).

**Key findings:** From the in vitro drug screening, we identified 21 drugs and tested 3 *in vivo* on the mdx mice. None of the three drugs significantly improved muscle histopathology.

**Significance:** The *in vitro* drug screen identified various efficient compounds, none of them strongly inhibited fibrosis in skeletal muscle of mdx mice. To explain these observations, we hypothesize that in Duchenne Muscular Dystrophy, in which fibrosis is a secondary event due to chronic degeneration and inflammation, the drugs tested could have adverse effect on regeneration or inflammation, balancing off any positive effects and leading to the absence of significant results.

## Introduction

Acute tissue injury generates transient inflammation and extracellular matrix (ECM) deposition which return to basal levels after the regenerative process is complete. However, under certain pathologic conditions, persistent damage and inflammation within the tissue generates excessive and chronic deposition of ECM components. This condition of persistent inflammation and elevated ECM is known as fibrosis [1,2]. Fibrosis hinder tissue regeneration and contributes to organ malfunction in different pathologies such as liver and kidney diseases, idiopathic pulmonary fibrosis, heart failure, and muscular dystrophies. Although fibrosis contributes to 45% of mortality in developed countries, the mechanisms regulating the initiation and the establishment of fibrosis have not yet been completely elucidated[3,4]. Despite the extensive study of fibrogenesis in response to injury, essentially no effective anti-fibrotic therapy is yet available. A better understanding of the cellular effectors and the molecular signals regulating this pathological condition is necessary.

In skeletal muscle, an organ with a high regeneration potential, fibrosis is a hallmark of severe muscular dystrophies. This is the case of the incurable Duchenne muscular dystrophy (DMD), where the lack of dystrophin protein leads to cycles of impaired regeneration, resulting in chronic degeneration of the tissue [5–7]. At the cellular level, fibroblasts are the master mediators of tissue fibrosis[8]. In the past 10 years, a group of tissue-resident multipotent mesenchymal progenitors have been described as precursors of fibroblasts. In adult muscles, these cells were named fibro/adipogenic progenitors (FAPs), based on their spontaneous potential to differentiate into myofibroblasts and adipocytes, both *in vivo* and in *vitro* [9–11]. Damage induces FAP activation and expansion, which are modulated by inflammatory signals, including Interleukin-4 and 13 (IL-4, IL-13), Tumor Necrosis Factor α (TNFα), and Transforming Growth Factor β (TGFβ) [9,12– 17]. Following acute damage, activated FAPs provide trophic support to satellite cells, required for efficient normal regeneration[18–21]. However, in degenerative condition such as in chronically damaged muscles of DMD patients, tissue clearance of FAPs fails. Chronic damage primes FAPs to differentiate towards both adipocytes and fibroblasts, leading to fibrofatty infiltration, and eventually loss of muscle function[15,22,23]. Due to their dual role in skeletal muscle regeneration/degeneration, FAPs are an ideal cellular target to improve regeneration and prevent fibrofatty deposition[19]. Thus, interventions targeting molecular pathways involved in FAP activation and differentiation are an attractive strategy to combat fibrotic diseases successfully.

Inhibition of the TGFβ signaling pathway during skeletal muscle regeneration reduces FAP number and down-regulates fibrillar collagen type 1 deposition, a hallmark of fibrosis [10,15,24,25]. TGFβ plays an essential role in tissue modeling and remodeling, and therefore, it is considered a master molecule in the initiation and establishment of fibrosis[26]. The canonical TGFβ pathway classically transmits extracellular signals via transmembrane serine/threonine kinase receptors and intracellularly via SMAD2/3/4 proteins. The non-canonical TGFβ pathways include a variety of intracellular cascades activated by TGFβ, independent of SMAD2/3/4. These include the molecules TGFβ-activated kinase 1 (TAK1), mitogen-activated protein kinase (MAPK) such as P38, extracellular signal-regulated kinases (ERK), and JUN N-terminal kinase (JNK), as well as nuclear factor kappa-light chain enhancer of activated B cell (NFκB), among others[27,28]. Therapeutic manipulation of the canonical and non-canonical TGFβ pathways have been shown to be beneficial in multiple myopathic states and fibrosis of various tissues[29].

Here, we used the Collagen1a1*3.6 EGFP mice to isolate and culture FAPs and analyze their differentiation into collagen producing cells, by following the expression of the EGFP reporter[30]. We performed a drug screening on freshly isolated FAPs, which allowed us to investigate the role of critical signalling pathways in regulating their fibrogenic differentiation.

Mdx mice (mouse model for the DMD) were treated with compounds that were found to strongly inhibit Collagen expression in vitro. However, none of the drugs reduced fibrosis in vivo. Here, we report that using an *in vitro* drug screening approach is a feasible technique to delineate pathways leading to the activation of a fibrogenic program. However, in the context of the DMD fibrosis cannot be specifically targeted independently of the other disease components: inflammation and muscle regeneration.

## Results

### 1. Collagen1a1*3.6 EGFP transgenic mice is a reliable and powerful tool for analyzing collagen expression induced by TGFβ1

We implemented an *in vitro* model to test compounds for the ability to block or interfere with FAP differentiation along the fibrogenic lineage. Knowing that increased collagen type 1 expression and deposition is a major hallmark of fibrosis [31], we took advantage of the Collagen1a1*3.6 EGFP transgenic mouse[30]. These mice express the EGFP gene under the control of a 3.6 kb fragment of rat procollagen type 1 alpha 1 (*Col1a1*) regulatory sequences (upstream promoter sequence). The use of this model enables the direct identification of any cell type that actively expresses high levels of *Col1a1* type 1 based on EGFP fluorescence.

To establish a suitable model capable of tracking FAP differentiation *in vitro*, EGFP negative FAPs were isolated from *Tibialis Anterior* (TA) of the Collagen1a1-3.6*EGFP mice 3 days after NTX damage and placed in culture[15,32]. Once they reached 60% confluency, FAPs were treated with 1ng.ml^-1^ of TGFβ1 for 72hr, and the percentage of EGFP positive cells was quantified using flow cytometry (Figure 1A). As suggested by previous studies[15], the physiological concentration of 1ng.ml^-1^ was sufficient to increase EGFP expression by 350% (Figure 1B and C) These results confirm that FAPs from the Collagen1a1*3.6 EGFP transgenic mice can be used as a robust *in vitro* model for screening potential blockers of *Col1a1* expression.

**Figure 1:**
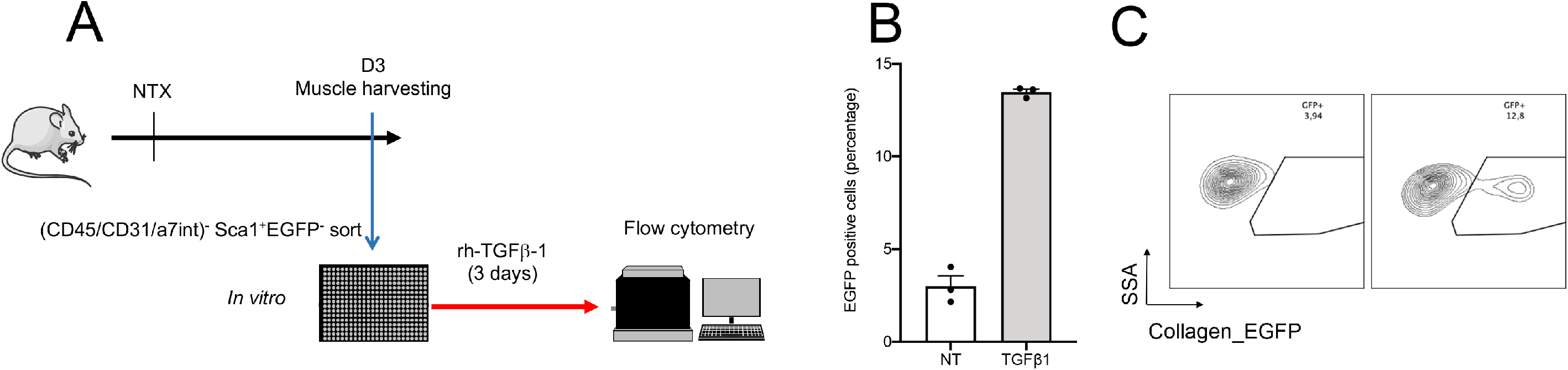
Collagen1a1*3.6 EGFP transgenic mouse is a powerful tool for analyzing TGFβ1 signaling. (**A**) Collagen1a1*3.6 EGFP mice were injected with notexin (NTX) in the tibialis anterior (TA) muscle at day 0 (D0). 3 days post injury, Collagen-EGFP negative FAPs were cell sorted and plated before been treated with 1 ng/ml of TGFβ1 for 72 hours (**B-C**). GFP expression was analyzed by flow cytometry as a measure of Collagen gene expression. n = 3 biological replicates. Treatment versus NT: p<0.01, ***

### 2. Drug screening in FAPs reveals potential targets for muscle fibrosis therapy

To evaluate the specific molecular pathways regulating FAP fibrogenic differentiation, we performed a drug screening using the *in vitro* model previously described and validated (Figure 1). To do so, EGFP negative FAPs were sorted from TAs and treated with 1 ng.ml^-1^ of TGFβ1 alone or with a set of 722 compounds organized into two libraries, a tyrosine kinase inhibitor library (KIL, 481 compounds) and a TOOL compound library (TOOL, 241 compounds) (Figure S1A, Table 1, and Table 2). Then, the percentage of EGFP positive cells was quantified by using a Cellomics Array scan (Figure S1A). Non-toxic compounds (see methods) with the ability to reduce the expression TGFβ1-induced EGFP expression by 50% and with a p-value < 0.05 were selected as positive hits (Tables 1 and 2 Figure 2A, B and S1A). Although 141 compounds met all the selection criteria, only 60 were selected for further assessment as these were compounds available in the market or in advanced phases of clinical development (Tables 1 and 2). For the second round of screening, EGFP negative FAPs were treated as previously described with the selected compounds, and the percentage EGFP positive cells was quantified by flow cytometry (Figure S1A). Figures 2C and 2E show the 21 most potent hits of both libraries that were then further tested in dose-response experiments. A total of 21 compounds showed a dose-dependent decrease in the percentage of EGFP positive cells induced by TGFβ1 (p<0.05, Figure 2C and E, Table 3). In the KIL library, MAPK inhibitors (VX702 and VX745), tyrosine kinase inhibitors (TyrK) (Nilotinib and Sorafenib), and the Farnesyltransferase inhibitor Tipifarnib significantly reduced the percentage of EGFP positive cells (p<0.05 or below, Figure 2C). In the TOOL library, the Bromodomain inhibitor (BRDI) I-BET151, the histone deacetylase (HDAC) inhibitor PCI-24781 and the PKC activator Indolactam V produced statistically significant reductions in EGFP positive cells (p<0.05 or below, Figure 2E). The most potent hits, with a translational potential from both libraries, were further tested at different doses and dose-response relationships were established (0.1-5 μM) (Figure 2D and F). These results show that targets of the TyrK inhibitor and the BRDI families regulate the establishment or maintenance of collagen expression and therefore could be investigated as potential therapies for fibrotic diseases.

**Figure 2:**
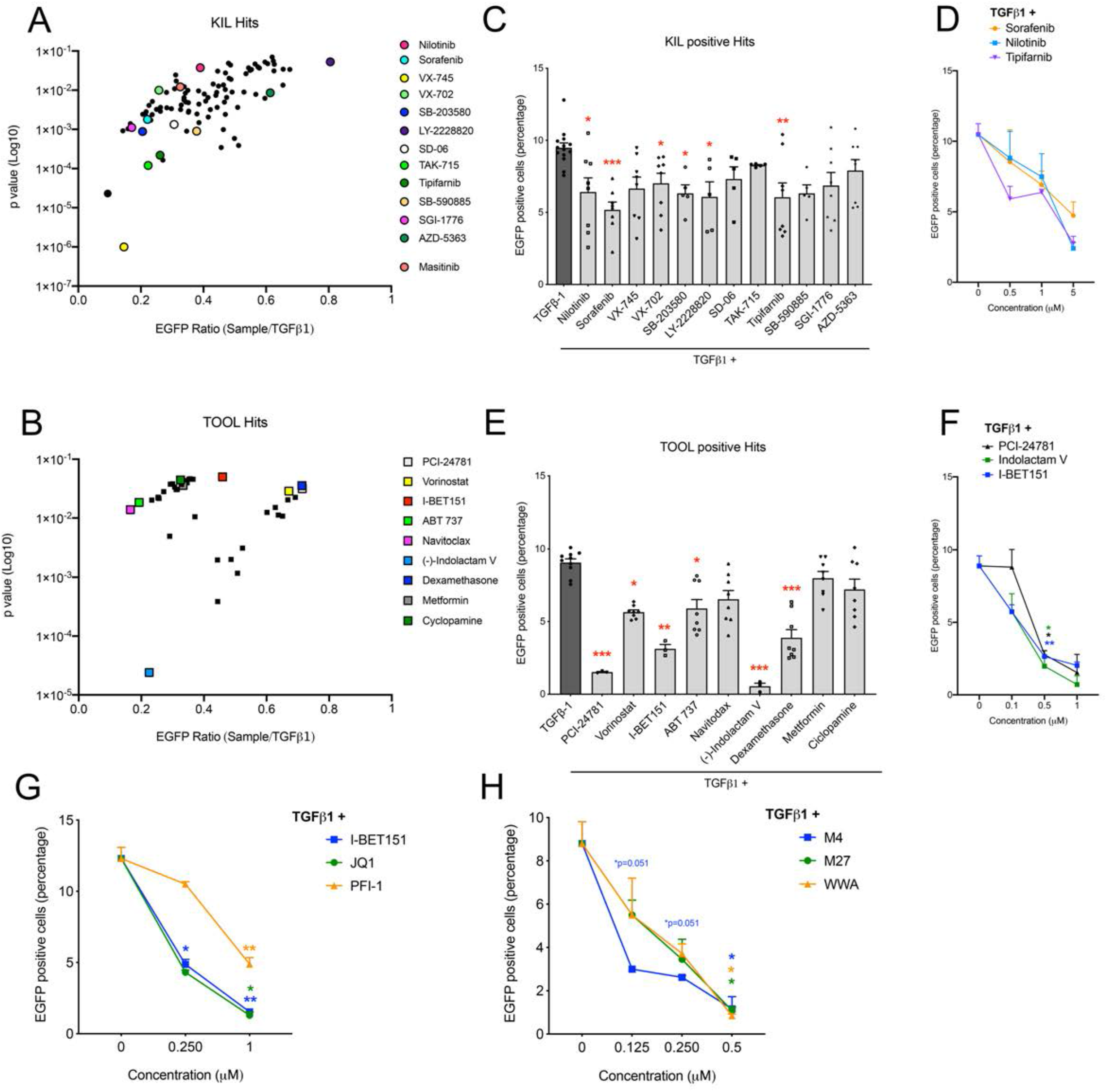
Drug screening targeting Collagen type I expression. (**A-B**) Tibialis anterior (TA) muscles of Collagen1a1*3.6 EGFP mice were injected with notexin (NTX). Three days after injury, EGFP negative FAPs were cell sorted and treated with TGFβ1 alone or with compounds from the KIL and TOOL libraries. Collagen-GFP expression was quantified using a Cellomic Arrayscan (*Figure S1*). 4 to 9 experimental replicates were performed (**C**) Sorted EGFP negative FAPs were treated with TGFβ1 and with selected compounds from the first round of screening of the KIL library. Percentage of EGFP positive cells was quantified by flow cytometry. Each technical replicate is presented performed (5 to 14) (**D**) Selected positive hits from the KIL library (Sorafenib, Nilotinib and Tipifarnib) were then tested as described at concentrations ranging from 0.5 to 5 μM. 2 biological replicates were performed. (**E**) Sorted EGFP negative FAPs were treated with TGFβ1 and with selected compounds from the first round of screening of the TOOL library. GFP expression was quantified using flow cytometry. Each technical replicate is presented performed (3 to 10) (**F**) Selected compounds from the TOOL library (Indolactam V, I-BET 151 and PCI-24782) were then tested in a dose dependent manner from 0.5 to 1 μM. 2 to 4 biological replicates were performed (**G**) Compounds from the “epiprobe” library were tested at 0.25 and 1 μM. Collagen-GFP expression was quantified using flow cytometry. 3 biological replicates were performed Drug versus TGFβ1: *: p <0.05, **: p<0.01

Interestingly, epigenetic modulators such as the BRDI I-BET151, and the HDAC inhibitor PCI-24781, were among the strongest inhibitors of *Col1a1* induction in our screen (as evidenced by EGFP expression)[33]. Based on these results, we tested a drug library composed of epigenetic regulators was tested by following the same procedure described above (Figure S1A). Among 26 epigenetic regulators (Table 4), the only group of compounds able to reduce the fluorescence induced by TGFβ1 were the members of the BRDI family: JQ1, and PFI-1 (Figure 2G). As already demonstrated in the heart, we confirm that the BRDI family members are essential regulators of FAP differentiation into fibroblasts [34]

Several transcriptional modifiers and regulators have been described as final effectors of the TGFβ signaling as well as mediating its involvement in fibroblast activation and scar deposition. These include serum responsive factor (SRF)[35], C-ets-1 (ETS1)[36], and NFkB[37]. NFkB pathway modulation has been associated with improvement in muscle health in mdx mice[38–40]. These results suggest that NFκB might be modulating FAP fibrogenic differentiation and warranted further screening for non-toxic NFκB inhibitors such as Withaferin[41]. Thus, we tested the ability of this compound to reduce collagen expression *in vitro* by using our previously described method (Figure S1A). Withaferin (WWA) and two of its synthetic analogues (M4 and M27) reduced the percentage of EGFP positive FAPs *in vitro* at a 0.5 μM dose (respectively -87%, -88%, and -90%, p<0.05) (Figure 2H). Overall, these results suggest that FAP differentiation into collagen-producing cells can be regulated by multiple signaling pathways, some of which could be independent of the canonical TGFβ signaling pathway. These results add layers of complexity and suggest that compensatory pathways could influence the results while testing drugs *in vivo*.

### 3. Non-canonical TGFβ pathway regulates FAPs fibrogenic differentiation

To validate that the candidate drugs act by another signaling than inhibiting the TGFβ pathway, we measured the level of activation of the downstream protein p38 MAPK (p-p38) in C3H10T1/2 mesenchymal progenitor cells (Figure 3A and B). As previously described[12,15], the treatment with TGFβ1 increased the phosphorylation levels of p38 by 2.48-fold. Interestingly, while Nilotinib and Sorafenib are known as TyrK inhibitors, co-treatment of C3H10T1/2 cells with TGFβ1 and these inhibitors blocked p38 phosphorylation, suggesting that their action works through the p38 MAPK signaling pathway (Figure 3B, respectively -56% and -59%). On the other hand, Masitinib, another well-known TyrK inhibitor (targeting Stem Cell Factor (SCF), c-Kit or PDGFRα) did not reduce p38 phosphorylation in our system. One of the possibilities is that the dose used in this assay was too low to induce any effect, as a decrease in EGFP positive FAP is noticeable only at 5μM for Masitinib (Figure S2). Lastly, while knocking-out BRD4 or using JQ1 *in vitro* on LPS-stimulated microglia is known to induce dephosphorylation of p38[42], TGFβ-induced p38 MAPK phosphorylation was not affected by JQ1 in our system (Figure 3A and B).

**Figure 3:**
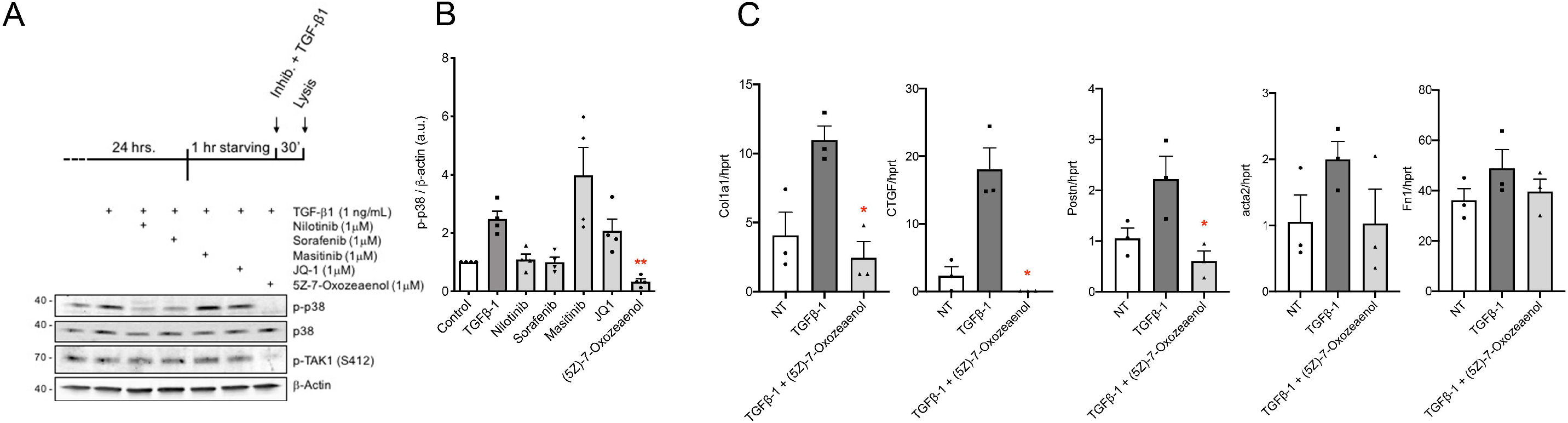
TAK1-p38 axis regulates FAP differentiation into fibroblasts. (**A-B**) C3H10T1/2 cell line was stimulated for 30 min with rh-TGFβ1 (1 ng/ml) alone or with 1 μM of Nilotinib, Sorafenib, Masitinib, JQ-1, or (5Z)-7-Oxozeanol. Protein lysates were extracted and Western Blot for p38, phospho-p38, phospho-TAK1 and β-Actin performed, and the data quantified. n=4. (**C**) EGFP negative FAPs were treated for 6 hours with 1 ng/ml of rh-TGFβ1 with or without 1 uM of (5Z)-7-Oxozeanol. Gene expression of *Col1a1, Acta2, CTGF, Fn1*, and *Postn* was quantified by digital droplet PCR and normalized to the expression of *hprt*. n=3. Treatment + rh-TGFβ1 versus rh-TGFβ1: *: p<0.05; **: p<0.01

In the context of fibrosis, the importance of the SMAD-independent TGFβ signaling pathway has been highlighted in various fibrosis models and the involvement of TAK1, a common upstream component of this pathway, as a key regulator of fibrosis has been shown in several tissues including kidney[43], skin[44], liver[45], and heart[46,47]. In muscle, TAK1 has been implicated in the regulation of MuSC fate and myofiber maintenance[48–50]. Moreover, TAK1 is a known activator of the p38 MAPK pathway[51–53], and indeed, the use of a pharmacological inhibitor (5Z-7-Oxozeaenol) decreased TGFβ1-induced phosphorylation of p38 MAPK (p<0.01; Figure 3A and B). To evaluate the participation of TAK1 in FAP differentiation towards fibroblasts, we evaluated the expression of fibrogenic genes by muscle FAPs *in vitro*. Sorted FAPs were treated with TGFβ1 (1ng.ml^-1^) with or without 5z-7-Oxozeaenol (1μM) for 6 hours (Figure 3C). In accordance with our previous results, the increase in gene expression of Collagen 1a1 (*Col1a1*), connective tissue growth factor (*CTGF*), periostin (*Postn*), Smooth muscle actin (*Acta2*), and fibronectin (*Fn1*) induced by TGFβ1 was attenuated in FAPs incubated with the TAK1 inhibitor 5Z-7-Oxozeaenol (p<0.05 or bellow, Figure 3C).

Taken together, these results suggest that non-canonical TGFβ signaling pathways participate in regulating the fibrogenic differentiation of FAPs. Also, we established that central components of this pathways such as TAK1 are playing an important role in modulating FAP fate, which may have significant translational potential for future clinical use.

### 4. Testing anti-fibrotic drugs candidates in vivo

To further validate *in vivo* our *in vitro* screening results, mdx mice (a murine model for DMD) were fed with a diet containing JQ1 from 4-5-weeks of age for one year (Figure 4A). Overall, the growth rate of treated animals was similar to that of untreated controls, suggesting the absence of systemic toxicity (Figure S3A). However, we noticed that mice fed with JQ1 display muscle mass loss specifically in the Quadriceps (−18%, p<0.09; Figure S3B). While collagen deposition (Picrosirius Red coloration (PSR)) in the diaphragm was not affected by the JQ1 diet, the size of myofibers was strongly decreased (−23%, p<0.05, Figure 4B to F). JQ1 has previously been demonstrated to have antifibrotic activity in the heart in both aortic constriction and post-myocardial infarction settings [34,54], as well as in bleomycin-injured lungs [55]. A possible reason for the discrepancy in the results we obtained from skeletal muscle is that while the studies describing the effect of JQ1 in the heart involved acute injury models, the model we used in this study is a chronic fibrotic disease with a strong inflammatory component. One alternative possibility is that our experimental animal models may not be representative of the clinical picture in patients with fibrosis. The mdx mouse is the most widely used murine model of DMD; however, it does not entirely recapitulate the fibro-fatty progression observed in humans. Indeed, mdx mice develop less fibrosis and with a later onset compared to human patients. In addition, in the murine model the fibrosis is primarily confined to the diaphragm muscle[56]. Other murine models, such as the mdx:utr^+/-^, which also lacks one allele of utrophin, a functional analog of dystrophin, have been proposed as a better alternative [57,58]. Indeed, mdx:utr^+/-^ mice mimic the human disease more closely and develop more severe fibrosis earlier after disease onset, including in the limb musculature [59,60]. As a result, we took advantage of the mdx:utr^+/-^ mouse model to further test the effect of JQ1 on muscle fibrosis. We performed continuous infusion of JQ1 (30mg/kg/day) by implanting mice with subcutaneous osmotic minipumps for 4 weeks (5 to 9-weeks-old, Figure 4G). The continuous delivery of JQ1 reduced post-natal growth, with 9-weeks-old mouse weight at decreased by 10% compared to the control group (p<0.05, Figure S3C). Concomitant to this, muscle mass of both gastrocnemius and quadriceps were reduced (respectively -11% p<0.01 and-11.3% p<0.001) (Figure S3D). Moreover, while the diaphragm’s histopathology was not affected (Figure 4H to L), collagen deposition of TA and gastrocnemius was decreased by 15% and 10% (p<0.5; Figure S3E to G). However, this was not associated with changes in myofiber size (Figure S3H to J).

**Figure 4:**
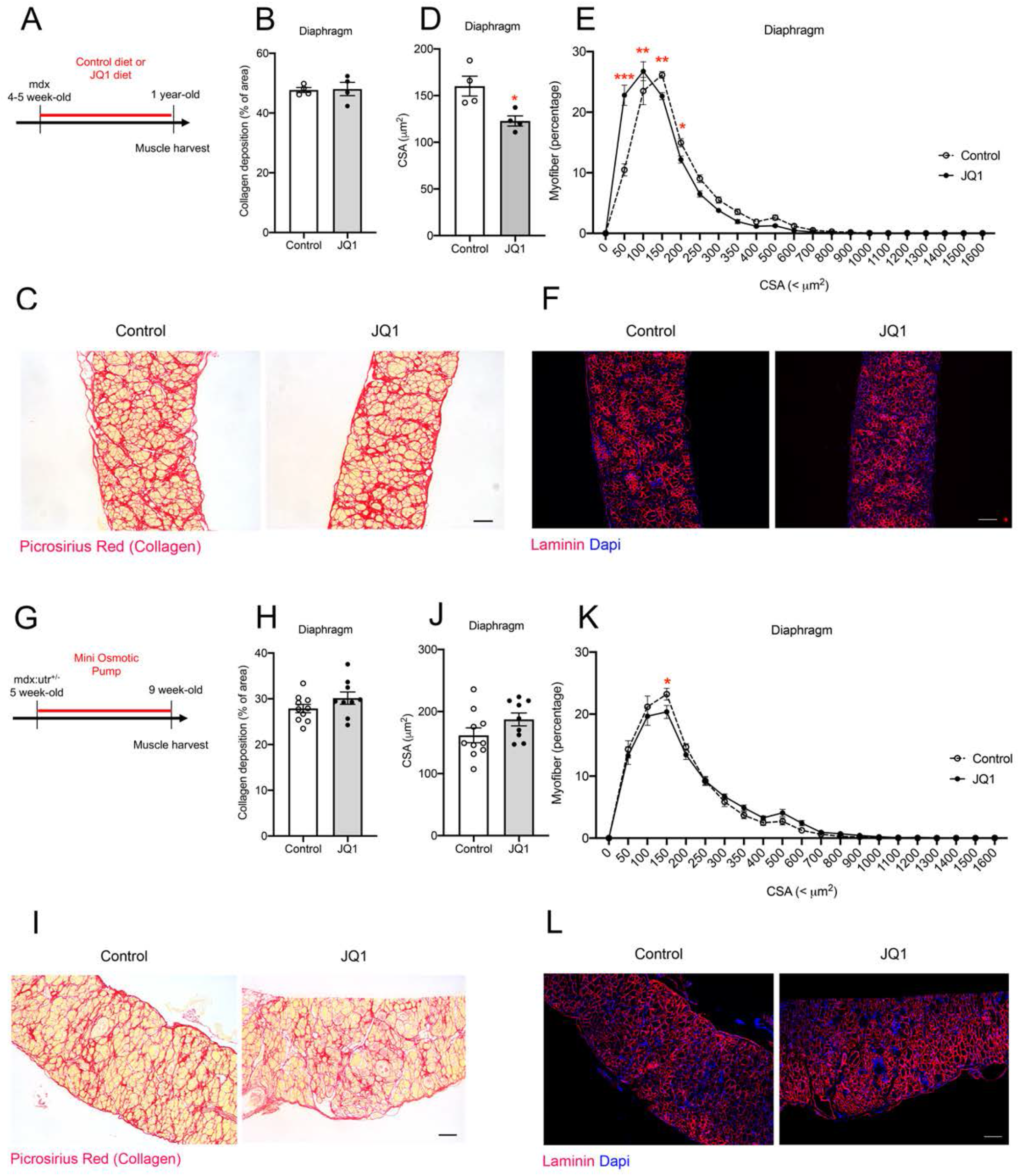
Treatment of JQ1 worsens DMD pathology. (**A-F**) mdx mice were fed with a control or a JQ1-medicated diet (30 mg/kg/day) for up to a year. Diaphragm collagen deposition (**B-C**) and myofiber size (**D-F**) were quantified. n=4 (**G-L**) mdx:utr^+/-^ mice were implanted with an Alzet osmotic pumps containing JQ1 or its vehicle for 4 weeks. Diaphragm collagen deposition (**H-I**) and myofiber size (**J-L**) were quantified. n=10 to 12 JQ1 versus control: * p<0.05; ** p<0.01; *** p<0.001 Scale bar = 100 μm

We then decided to test the TyrK inhibitors Nilotinib and Sorafenib. While we previously demonstrated that Nilotinib strongly inhibits fibrosis in the muscle and the heart after acute damage[15,61], Nilotinib administration has been associated with adverse effects such as hyperglycemia, increased LDL and HDL, vascular and cardiovascular toxicity when used in the context of long-term treatment[62]. Very recently, White et al., demonstrated that DMD and Becker muscular dystrophy (BMD) patients displayed plasma lipid abnormalities early in the onset of the disease[63]. In addition, *in vitro*, Nilotinib inhibits C2C12 myoblast differentiation[64]. Thus, Nilotinib is unlikely to be viable as drug for long-term treatment of DMD patients.

In contrast, Masitinib, which is active on a similar range of substrates, does not demonstrate toxicity when tested *in vivo* and *in vitro*[65]. To note, while Masitinib demonstrated great inhibitory activity during our first screening, it did not pass our second screening as it was notable to decrease the percentage of EGFP positive cells at concentrations in the therapeutic range (Figure 2A and S2). Masitinib was injected i.p. every day for two weeks in mdx:utr^+/-^ mice at 60 mg/kg/day (Figure 5A). No differences in mouse weight were noticed, confirming the absence of toxicity (Figure S4A). However, only a slight decrease in total collagen content was detected by PSR in the diaphragm (−12.8%, p<0.05; Figure 5B and C, Figure S4B). This was associated with no changes in myofiber size in the diaphragm, while a 16% decrease was observed in the TA (p<0.05; Figure 5D to F and S4C). Finally, to determine whether the results we observed were influenced by to the route of drug administration, we delivered Masitinib for 8 weeks by osmotic minipump implantation (Figure 5G). However, despite this method, no improvement of fibrosis deposition and muscle histopathology was observed (Figure 5H, I, J and N, Figure S4D to F). Lastly, mice were treated with Sorafenib, another promising TyRK inhibitor similar to nilotinib, by using the osmotic pump delivery method for 8 weeks (Figure 5G). Overall, no differences were noticeable on mouse body weight, muscle mass, collagen deposition and myofiber size (Figure 5H, K, L, M and N; Figure S4G to I).

**Figure 5:**
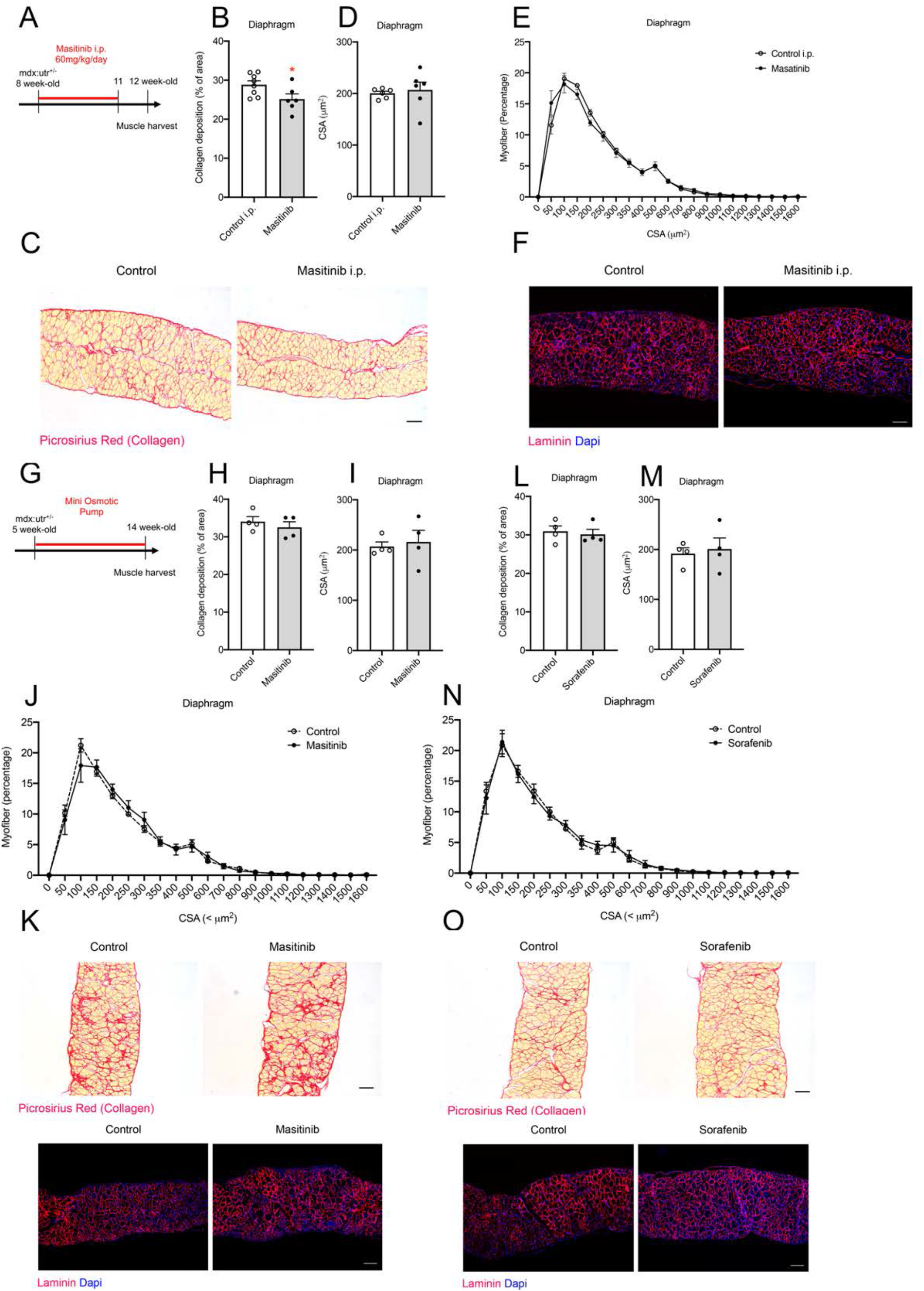
Sorafenib and Masitinib treatments do not improve DMD histopathology. (**A-F**) mdx:utr^+/-^ mice were injected i.p. with 60 mg/day/kg of Masitinib from 8 to 11-weeks-old. Diaphragm collagen deposition (**B-C**) and myofiber size (**D-F**) were quantified. n=6 to 8 (**G-O**) mdx:utr^+/-^ mice were implanted with osmotic pumps containing either Masitinib, Sorafenib or the appropriate control vehicle from 5 to 14-weeks-old. Diaphragm collagen deposition (**H, L, K, O**) and myofiber size (**I, J, M, N**) were quantified. n=4 Treated mice versus control: *:p <0.05, Scale bar = 100 um

## Discussion

There is currently no cure for fibrosis, mainly because this condition is a multifactorial factors and likely multiple molecular pathways are involved in triggering, establishing, and maintaining scar-forming disorders and related pathologies. Therapeutic strategies to reduce fibrosis in different tissues have been attempted with limited success [2]. Most of the information that we know about the fate of FAPs and their contribution to muscle regeneration and repair has been done by using skeletal muscle as a model to study fibrosis [9–12,15,20,21,23,66]. Here, we established a screening system using primary FAP cell culture to: 1) Define the intracellular pathways leading to the activation of a fibrogenic program in response to TGFβ1; 2) Identify a set of drugs capable of interfering with the fibrogenic differentiation of FAPs, to test their therapeutic potential. We found that compounds such as Nilotinib, Sorafenib, JQ1, I-BET151, and Withaferin were able to decrease the expression of Collagen type 1 induced by TGFβ1.

Using muscle resident FAPs, we first confirmed that TGFβ1 upregulates the expression of *Col1a1*. (Figure 1A-C). To note, collagen expression might not be representative of the other ECM genes such as CTGF/CCN2 or fibronectin, which actively participate in the installation of fibrosis[67,68]. We showed that the local rise of TGFβ concentration in skeletal muscle after acute damage pushes FAPs towards fibrogenic differentiation. Similar results have been obtained by others and ourselves, where the expansion of resident FAPs is closely associated with elevated TGFβ levels and increased ECM deposition during regeneration and repair[22,69,70]. In addition, we previously showed a temporal correlation between the peak of TGFβ expression in the tissue and the *Col1a1* levels after skeletal muscle acute damage at day 7 [15,20].

For some of the compounds emerging form our screen, an anti-fibrotic effect had previously been demonstrated in different organs, including lung, kidney, liver, heart, and skeletal muscle. Members of the TyrK inhibitor family such as Imatinib[25,71,72], Nilotinib[15,20,73–75], Sorafenib[76], Sunitinib[77]; and epigenetic regulators such as BRD4 and HDAC inhibitors have been effective in blocking fibrosis[55,78–81]. BRD regulators specifically recognize acetylated lysine residues, which act as scaffolds and attract components of the transcriptional machinery to the acetylated lysine residues of histones, resulting in modulation of gene transcription. This finding is particularly interesting because the role(s) of epigenetic regulators in fibrosis and their therapeutic potential is still poorly understood[82]. Pharmacologic modulation of epigenetic readers is emerging as a novel therapeutic approach for the treatment of inflammatory diseases[82].

Here, we identify FAPs as pharmacological targets for the action of TyrK, BRD and p38 MAPK inhibitors. The TGFβ–TAK1 pathway is a shared molecular pathway regulating TyrK, BRD and p38 MAPK signaling. Growing evidence supports the role of TAK1 as a significant regulator of TGFβ signaling through the regulation of the profibrotic response in several systems[43–45,83– 87]. Very recently, it has been shown that the action of Catalpol (an anti-inflammatory and antioxidant drug from Chinese medicinal herb *Rehmannia*) on DMD histopathology was due to its binding to TAK1[88,89]. Besides its role in fibrosis, TAK1 also has been described as an important mediator of carcinogenesis and cell survival[90]. In all these settings, TAK1 regulates inducible transcription factors such as NFkB, and other kinases such as p38 MAPK and c-Jun N terminal kinases (JNKs)[38,51–53,91]. In our initial drug screening, all the TAK1 inhibitors tested were found to be toxic, thus confirming its central role in cell survival. For this reason, we did not pursue the use of the TAK1 inhibitors *in vivo*.

We also tested a natural compound, Withaferin A, and two chemical derivatives, known for their inhibitory effect on the transcription factor NFkB[92,93]. Using the Col1a1-EGFP model presented here, we observed a decrease in EGFP positive cells in Withaferin A treated FAPs, suggesting that the NFkB factor might be playing a role in regulating the expression of collagen in FAPs. Similar results were found in the heart, where Withaferin A reduced type I collagen expression *in vitro* and inhibited the development of myocardial fibrosis *in vivo*[94]. Dysregulation of NFkB activity can contribute to chronic inflammatory diseases such as dystrophies. Treatments with NFkB inhibitors improved muscle function in mdx mice[39,40]. The use of Withaferin A has been associated with decreased inflammation in several models[95], and may be due to an effect on early inflammatory stages of fibrosis. However, in our hands, mice treated with Withaferin in i.p (IMS-088). displayed swelling of the abdominal area and peritoneal adhesions, resulting in the interruption of its testing *in vivo* (data not shown).

JQ1 is an inhibitor or the BET family of proteins which includes BRD2, 3 and 4[96]. Its action as an antifibrotic has been demonstrated in various tissues, including the heart, lungs, kidneys, and the liver[34,54,78,97,98]. In this study, we tested the action of JQ1 on muscle fibrosis with long-term treatment (medicated food for one year) and a short-term treatment (osmotic minipump for 4 weeks). While we did detect a small decrease in fibrosis in TA and GC with the short-term treatment, we also detected a substantial decrease in myofiber size in the diaphragm of the mice treated with JQ1 for a year, suggesting that the drug affects fiber maintenance or metabolism (Figure 4 and S3). We speculate that the absence of effect on matrix deposition commensurable with that observed *in vitro* may be due to an unbalance between the anti-fibrotic and the anti-inflammatory function of JQ1[99,100]. Lastly, the use of TyrK inhibitors, especially Nilotinib, *in vitro* and *in vivo* has shown promising results in various settings and fibrotic diseases[15,61,76,101,102]. However, Nilotinib has been shown to induce side effects in long-term treated patients[62], and off-target effects in skeletal muscle myogenic progenitors[64]. Because of that, and despite the fact that it reduces Col1a1-EGFP levels only at higher doses (5μM), we focused on the TyrR inhibitor Masitinib as a candidate to replace Nilotinib. Currently, Masitinib is used in 31 clinical trials (NIH clinicalTrials.gov). However, while it also induced a slight decrease in fibrosis content in the diaphragm, it did not improve the overall histopathology of the disease (Figure 5 and S4). Similar results were found with the TyrK inhibitor Sorafenib. One of the many possible explanations would be that the drugs, as does Nilotinib, also interact with immune cells, endothelial cells, and muscle cells (progenitors and myofibers)[64]. Indeed, MAPK p38, NFkB, KIT, PDGF, and FGF signaling pathways regulate immune cell functions[103,104], vascularization[105,106], or myogenic cell proliferation, differentiation, and myofiber growth[107–109].

In conclusion, here we show that in a model system in which fibrosis is a secondary event caused by dysfunction of the parenchymal muscle stem cells and fibers, inhibitors capable of preventing the activation of a fibrogenic transcriptional programme seem to be inefficient in protecting against the disease. These results contribute to the growing evidence that modeling fibrotic diseases *in vitro* to perform drug screening may not yield hits that will perform well in *in vivo* models[110,111].

## Materials and methods

### Animals

Mice were maintained in an enclosed and pathogen-free facility. Mice were housed in standard cages under 12 h light–dark cycles and fed *ad libitum* with a standard chow diet. All experimental procedures were approved by the University of British Columbia Animal Care Committee. Transgenic mice expressing the enhanced green fluorescent protein (EGFP) under a *Collagen1a1* enhancer, *Col1a1*3*.*6*-eGFP, were a gift from Pr. D.W. Rowe (Center for Regenerative Medicine and Skeletal Development, University of Connecticut Health Center, USA). mdx:utr^+/-^ mice were a gift from Dr. Lisa Hoffman (Western University, London, ON, Canada). Adult mice >8-weeks-old, both male and female, were used unless otherwise specified. Acute muscle damage was induced by intramuscular injection of 0.15 µg notexin (NTX) snake venom (Latoxan), into the tibialis anterior muscle (TA).

Mice were fed with control diet (Research Diet, #D111112201) or JQ1 diet (control diet supplemented with JQ1 at 25 mg/kg (Cayman Chemical #11187, or Selleckchem #S7110)).

Mice were daily treated by intraperitoneal (i.p.) injection with the vehicle (DMSO/ETOH=1:1), or Masitinib (ABSCIENCE, #AB1010, 60 mg/kg/day).

### Mini osmotic pump

The empty mini osmotic pump (ALZET; model 2002) was filled with the vehicle (DMSO/ETOH=1:1) or drug solution (JQ1: Cayman Chemical #11187 or Selleckchem #S711, 30 mg/kg/day; Masitinib: ABSCIENCE #AB1010, 60 mg/kg/day; Sorafenib: BAYER #BXA5X4R, 6 mg/kg/day) in sterile conditions. In order to allow the pump to equilibrate and reach its steady-state pumping rate, the filled pumps were primed overnight in sterile saline at 37°C. Animal were anesthetized and shaved. A 1 cm incision was performed on the skin and a pocket was created with 1 ml of sterile saline. The osmotic pump was slowly inserted, then the muscle layer was gently tented and closed with 6-0 monocryl. By using an 18G needle, a hole was then poked in the muscle layer. The catheter was fed through the hole until the first retention bead passed through and into the peritoneal space. To finish, the purse string suture was tightened using square knots. The catheter was allowed to sit naturally by adjusting the pump. Finally, the skin layer was closed using an intradermal pattern.

### Histology

Before tissue collection, animals were perfused transcardially with 20 ml of 1X PBS 4% PFA Tissues were processed for paraffin-embedding using standard methods. Sections of muscle tissues were stained with picrosirius red (PSR) in order to quantify collagen deposition.

For laminin staining, sections were deparaffinized and antigen retrieval performed in proteinase K buffer (Abcam, #ab64220) for 20 min at room temperature (RT). Slides were then washed in 1X PBS and then incubated in a blocking solution containing 3% normal goat serum and 0.3% triton X-100 in 1X PBS for 60 mins at RT prior to incubation with Laminin (Abcam #ab11575; 1:200) at 4°C overnight. Slides were then washed with 1X PBS and incubated in blocking solution containing the secondary antibodies (ThermoFisher) for 2 hours at RT. Following antibody incubation, 3 × 5 min PBS washes were performed, and sections were stained with DAPI for 10 min (ThermoFisher, #D3571, 0.6 μM) before being mounted with fluorescent mounting medium (Dako).

### Imaging

PSR and laminin images were acquired at 10X magnification using a Nikon Eclipse Ni equipped with a device camera (Nikon Digital Sight DS-U3 for brightfield, Qimaging Retiga EXi for fluorescence) and operated via NIS software. Collagen deposition and CSA were calculated using Fiji (ImageJ, version 2.0.0-rc/69/1.52n, NIH, MD) and Open-CSAM[112]. Images were assembled using Adobe Illustrator CS6 (Adobe)

### FAP culture

FAP were isolated and cultured as described in[32] with some modifications. Hindlimbs or damaged TAs were carefully dissected and gently torn with tissue forceps. Enzymatic digestion was performed with Collagenase D (Roche Biochemicals; 1.5 U/ml) and Dispase II (Roche Biochemicals; 2.4 U/ml), at 37 °C for 60 min. Preparations were passed through 70 µm and 40 µm cell strainers (Becton Dickenson), and washed in 1X PBS containing 2 mM EDTA and 2% FBS (FACS buffer). Resulting single cells were collected by centrifugation at 300g for 5 min. Cell homogenate was incubated with primary antibodies for 30 min at 4°C in FACS buffer. Monoclonal primary antibodies were used as following: anti-CD31 (eBioscience, clone: 390), anti-CD45 (AbLab, clone: I3/2), anti-Sca-1 (eBioscience, clone: D7) and anti-α7 integrin (AbLab, clone: R2F2). Cells were stained with Hoechst 33342 (2.5 µg.ml^-1^, Sigma) and resuspended in FACS buffer immediately before sorting. Sorting was performed on FACS Aria II (Becton Dickenson) or Influx (Becton Dickenson). Gates were defined based on fluorescence minus one (FMO) controls and EGFP negative FAPs were sorted as CD31/CD45/a7int-Sca1+ GFP-. For *in vitro* experiments, FAPs were seeded at a density of 10.000 cell/cm^2^ in high–glucose Dulbecco’s modified eagle medium (DMEM) (Invitrogen) supplemented with 10% FBS and 1.5 ng/ml β-FGF (Invitrogen).

### In vitro drug screening

EGFP negative FAPs were plated at 10,000 cells/cm^2^ in 384 well-plates (Falcon) and treated at 50-60% confluence, either with rh-TGFβ1 (1 ng.ml^-1^) (eBioscience) or rh-TGFβ1 + drugs in DMEM 5% FBS.

For the first screening, 6 to 8 mice were pooled before sort and four to nine technical replicates were performed. 722 chemical compounds (Table 1 and 2) organized into two libraries (KIL and TOOL, donated by Dr. Rima Al-awar (Ontario Cancer Institute, Toronto) were tested. After a 72h treatment period, cells were fixed with paraformaldehyde (PFA) 4% for 10 min, and Hoechst 33342 (Sigma) was used for nuclei staining. Viability and GFP expression were analyzed using Cellomics Array scan (Thermo Fisher Scientific).

For the second screening, 6 to 8 mice were pooled before sort, triplicate were performed and the experiment was repeated at least 2 times. screening was performed on 60 compounds in the same condition as previously described at a 1μM dose. After 72 hours, cells were detached from the plate using trypsin and resuspended in FACS buffer and Hoechst 33342 (Sigma). Percentage of EGFP positive cells was calculated on the total event that were Hoechst+. The most significant drugs were finally tested in a dose response (0.01 to 5 μM) and the percentage of FAP EGFP positive was analyzed by FACS.

An epigenetic library of 26 compounds was obtained from Chemical Probes and was screened by flow as previously described (concentrations of 0.25 and 1 μM). Withaferin and its derivatives compounds (Imstar Therapeutics) were tested at a range of 0.06 to 0.5 μM concentrations. Percentage of EGFP positive FAPs was quantified by flow cytometry as described above.

Lastly, once FAP reached 60% confluence, the cells were incubated in 5% FBS, no bFGF prior to being stimulated with rf-TGFβ1 (1 ng.ml^-1^) and later with or without 5-z-7Oxozeanol at 1 nM for 6 hours. RNA isolation was performed using RNAzol reagents as per the supplier’s instructions.

### RT-PCR and ddPCR

Reverse transcription was performed using 100 ng of RNA and Superscript Reverse Transcriptase according to the supplier’s instructions (Applied Biosystems). The cDNA was diluted ten times in

RNAse free water (Thermofisher) and 2.5 µl was used in a reaction mix containing Droplet Digital PCR Supermix (Bio-Rad), TaqMan assay and RNAse free water. The Taqman probes (Suppl Table 1) used were the following: droplets were generated with a QX100 droplet generator (Bio-Rad), after mixing 20 µl of reaction mix and 70 µl of droplet generator oil (Bio-Rad). The emulsified samples were loaded onto 96-well plates and endpoint PCRs were performed in C1000 Touch thermal cycler (Bio-Rad) at the following cycling conditions: 95 °C for 10 min, followed by 45 cycles at 94 °C for 30 s and 60 °C for 1 min, followed by 98 °C for 10 min. The droplets from each sample were read through the QX200 droplet reader (Bio-Rad). Resulting PCR-positive and PCR-negative droplets were counted using QuantaSoft software (Bio-Rad). Data for each gene were normalized to *hprt* expression.

### Cell culture

C3H10T1/2 is a cell line of mesenchymal progenitors (Clone 8, American Type Culture Collection, Manassas, USA), the cells need to be maintained at 50-70% confluence. C3H10T1/2 cells were grown at 37 °C in 5% CO2 in growth medium; high-glucose Dulbecco’s modified Eagle’s medium (DMEM) (Invitrogen), supplemented with 10% fetal bovine serum (FBS) and Penicillin/streptomycin. For western blot analysis, cells were serum-starved for 1 hr before being treated with rh-TGFβ-1 (e-Bioscience) in DMEM supplemented with 2% (v/v) FBS. Specific treatments the following inhibitors were also used at 1 μM final concentration: Nilotinib (Tasigna®, AMN107; Novartis), Sorafenib (Bayer), JQ-1 (Cayman Chemical), 5Z-7-Oxozeaenol (Cayman Chemical). Protein extracts were done after 30 min of incubation at 37 °C in 5% CO2.

### Protein extraction and western blot

Protein extracts from cells were obtained using RIPA 1X lysis buffer (#9806 Cell signaling, MA, USA) plus protease/phosphatase inhibitors (#P8340 and #P0044, Sigma-Aldrich, USA). Cells were sonicated for 10 s and centrifuged at 9000g. Proteins were quantified with the Micro BCA assay kit following the manufacturer’s instructions (Pierce, IL, USA). Extracts were subjected to SDS-PAGE electrophoresis in 9% polyacrylamide gels, transferred to PDVF membranes (Millipore, CA, USA), and probed with primary antibodies: rabbit anti-phospho-p38 (Thr180/Tyr182) (#9211S, Cell Signaling, USA), rabbit anti-p38 (#9212, Cell Signaling, USA) and mouse anti-β-actin (#21001901, AbLab, BC, CA). Then, primary antibodies were detected with IRDye® Infrared Dye labeled secondary antibodies (LI-COR). All immunoreactions were visualized by with an Odyssey® imaging system. Western blot densitometry quantification was done using Fiji (ImageJ) software. Protein levels were normalized with the levels of the loading control.

### Statistical analysis

Graph and statistical tests were performed using Prism 8 (GraphPad Software, La Jolla California, USA). Depending on the experiment, one-way or two-way ANOVA were performed, corrections were applied, followed by post-hoc test. Gaussian distribution was not assumed. A probability of <5% (p < 0.05) was considered statistically significant. Sample size and/or technical replicate number for each experiment is indicated in the figure legend. Graphs are represented as mean ± standard error of the mean. Figures were assembled using Adobe Illustrator CS6 (Adobe)

## Figure legends

***Figure S1: In vitro drug screening scheme***.

1^st^ screening:

Tibialis anterior (TA) muscles of Collagen1a1*3.6 EGFP mice were injected with notexin (NTX). Three days after injury, EGFP negative FAPs were sorted, plated in 384 well-plates and treated at 50-60% confluence, either with recombinant human TGFβ (1 ng.ml^-1^) (eBioscience) or TGFβ1 + drug. After 72 hours of treatment, cells were fixed and stained for nuclei detection. Viability and GFP expression were analyzed using Cellomics Array scan (Thermo Fisher Scientific).

2^nd^ screening:

EGFP negative FAPs were sorted from NTX-injected TAs, plated in 48 well-plates and treated when reached 50-60% confluence, either with rh-TGFβ (1 ng.ml^-1^) (eBioscience) or rh-TGFβ1 + drug. After 72 hours, cells were detached from the plate using trypsin, resuspended in FACS buffer and stained with Hoechst 33342. Percentage of EGFP positive cells was calculated based on the total Hoechst+ events.

***Figure S2: Masitinib inhibits Collagen-EGFP expression induced by TGFβ1***

Tibialis anterior (TA) muscles of Collagen1a1*3.6 EGFP mice were injected with notexin (NTX). Three days after injury, EGFP negative FAPs were cell sorted and plated before been treated with rh-TGFβ1 alone or with Masitinib as different doses (0.01 to 5μM. n= 1 to 2)

***Figure S3: In vivo validation of JQ1***

(**A-B**) mdx mice were fed with control diet or JQ1 diet (30 mg/kg/day) for up to a year. Mouse mass (**A**) and muscle mass (**B**) were measured. n=4

(**C-J**) mdx:utr^+/-^ mice were implanted with pumps containing JQ1 or its vehicle for 4 weeks. Mouse body mass (**C**) and muscle mass (**D**) were measured. TA, GC, and Quad collagen deposition (**E-G**), as well as myofiber size (**H-J**) were quantified. n=10 to 12

End-point versus starting point: $$$ p<0.001

JQ1 versus control: * p<0.05; ** p<0.01; *** p<0.001

***Figure S4: In vivo validation of Masitinib and Sorafenib***

(**A-C**) mdx:utr^+/-^ mice were injected in i.p. with Masitinib or its vehicle from 8 to 11-weeks-old. Mouse body mass was measured (**A**). TA, GC, and Quad collagen deposition (**B**), as well as myofiber size (**C**) were quantified. n=6 to 8

(**D-F**) mdx:utr^+/-^ mice were implanted with minipumps containing Masitinib or its vehicle for 9 weeks. Mouse body mass was measured (**D**). TA and GC collagen deposition (**E**), as well as myofiber size (**F**) were quantified. n=4

(**G-I**) mdx:utr^+/-^ mice were implanted with minipumps containing Sorafenib or its vehicle for 9 weeks. Mouse body mass was measured (**G**). TA and GC collagen deposition (**H**), as well as myofiber size (**I**) were quantified. n=4

End-point versus starting point: $ p<0.05; $$$ p<0.001

Treatment versus control: * p<0.05

## Acknowledgments

The authors would like to thank Professor David W. Rowe (Center for Regenerative Medicine and Skeletal Development, University of Connecticut Health Center, USA) and Doctor Lisa Hoffman (Western University, London, ON, Canada) for gifting us the Col1a1*3.6-eGFP and the mdx:utr+/-mouse colonies. We would like also to thank Dr. Rima Al-awar (Ontario Cancer Institute, Toronto) for kindly gifting us the TOOL and KIL libraries, as well as Andrew Johnson, Justin Wong (UBC Flow core), and Josh Hashimoto for their help.

## Author contributions

M.T., M.L., C.C., L.R., F.L., O.C., L.W.T., and A.W. were responsible for performing and analyzing experiments. M.T., M.L., H.S., and F.M.V.R. were involved in experimental design, data interpretation, and preparation of the manuscript. All authors were involved in editing the manuscript.

## Competing interests

The authors declare no competing or financial interests.

## Funding

This work was supported by the Fondation pour la Recherche Médicale (FRM; 40248 to M.T.); European Molecular Biology Organization (EMBO; ALTF 115-2016 to M.T.), by the Association contre les myopathies (AFM; 22576 to MT), by Michael Smith Foundation for Health Research (MSFHR; 18351 to MT), by the Agencia Nacional de Investigación y Desarrollo (ANID; POSTDOCTORADO BECAS CHILE/2011-74120052 to M.L.), by the Canadian Institutes of Health Research (CIHR-FDN-159908 to F.R.) and by the US Department of Defense (W81XWH-16-1-0327).

---

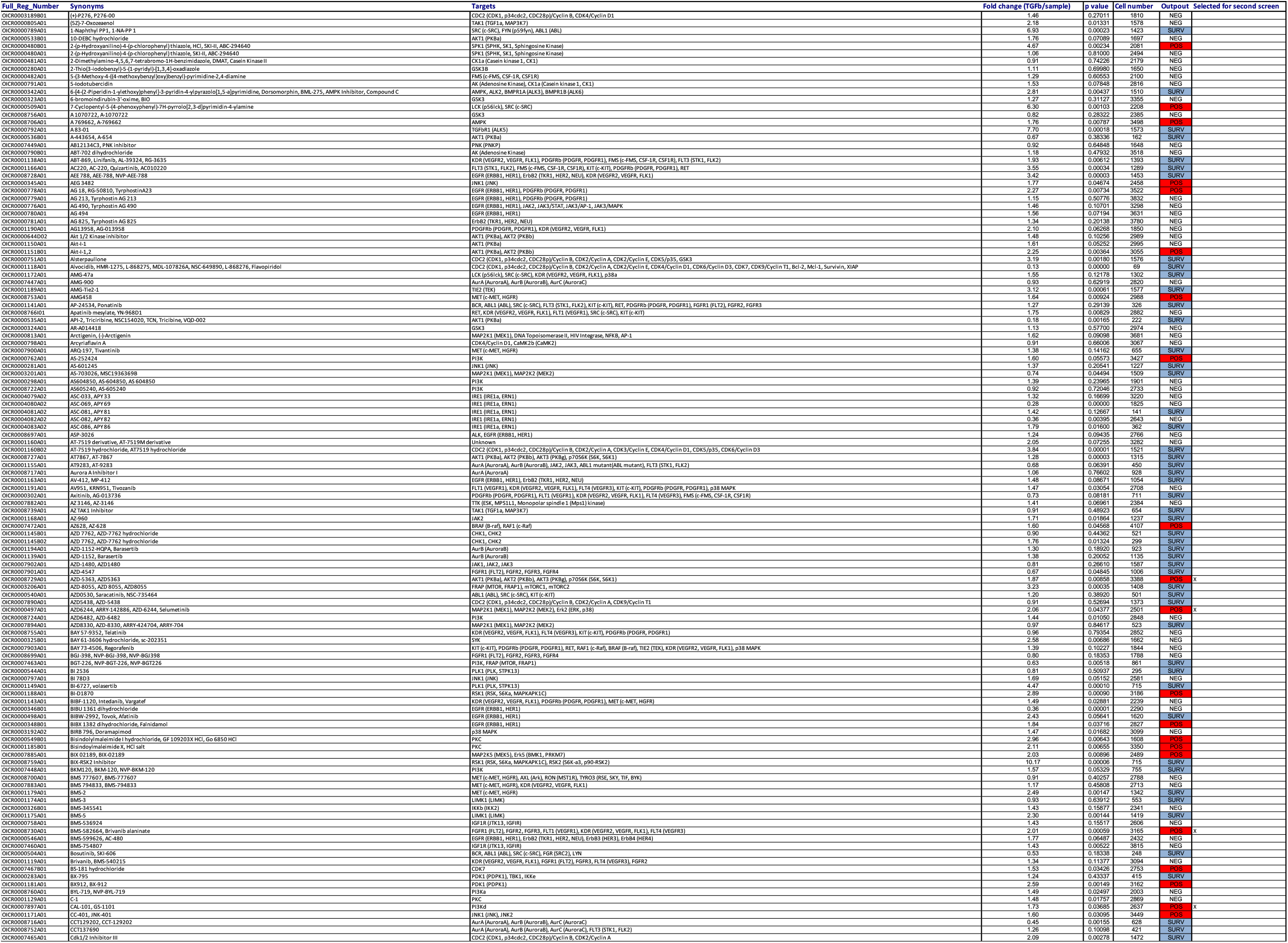
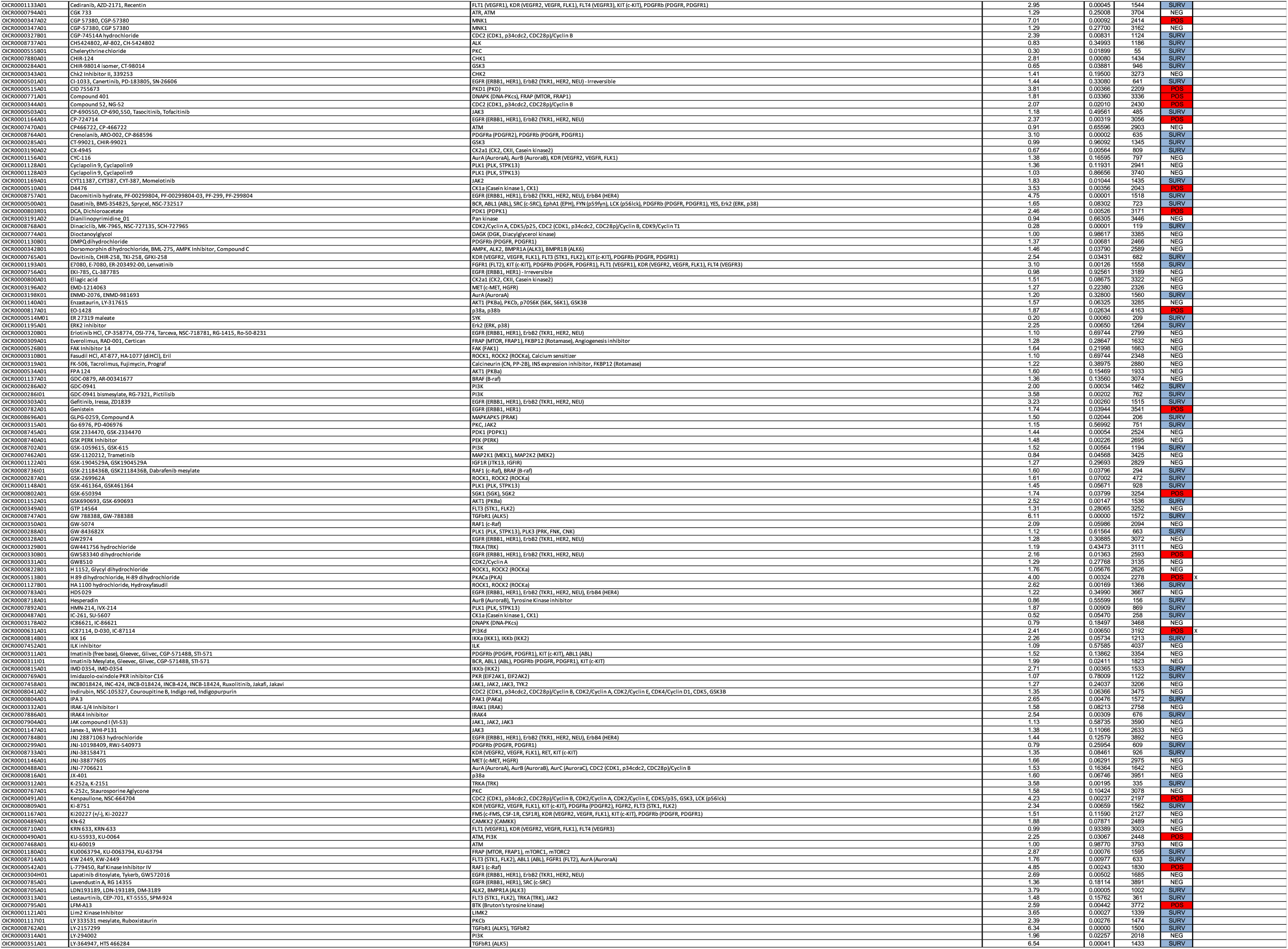
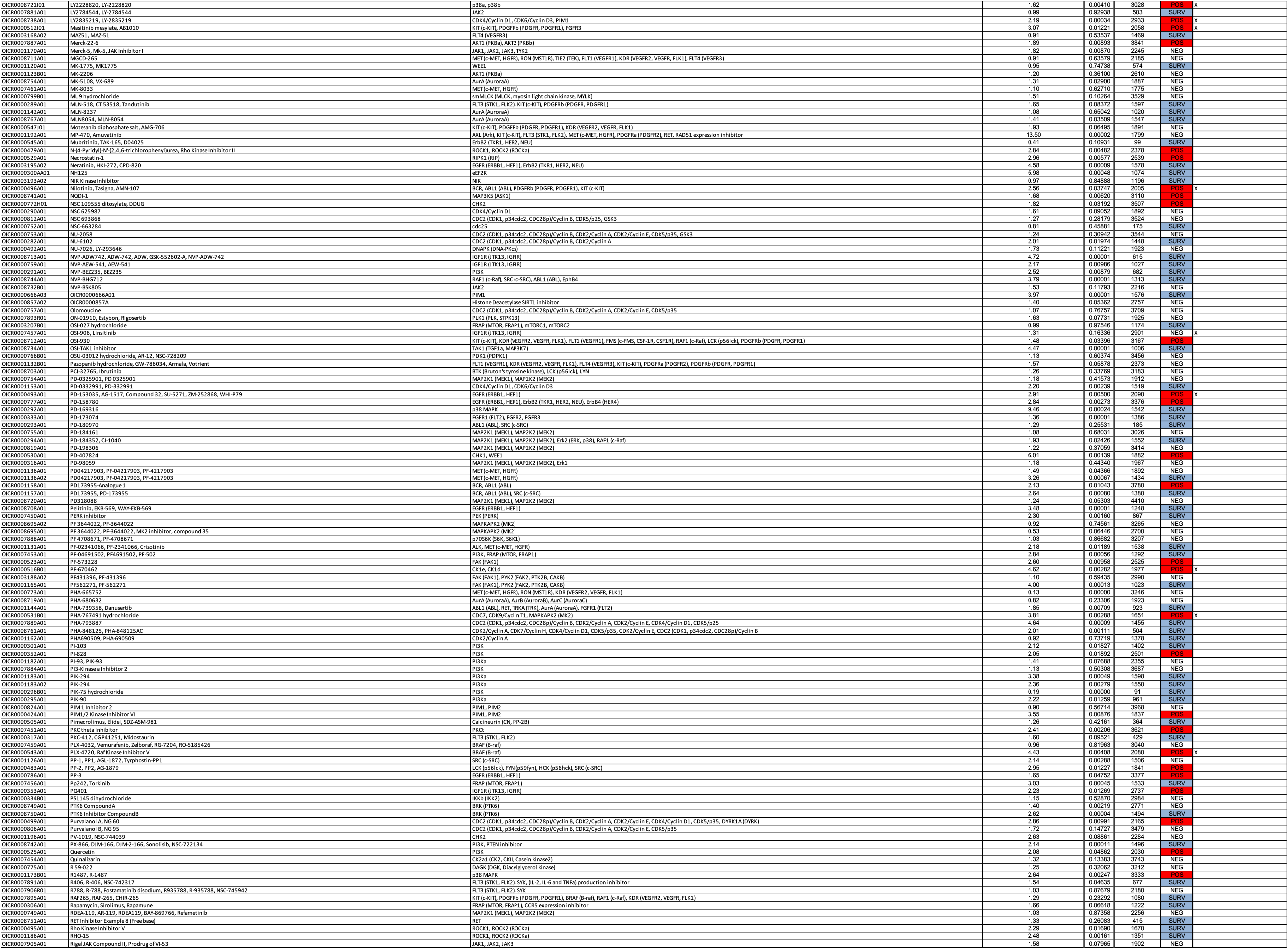
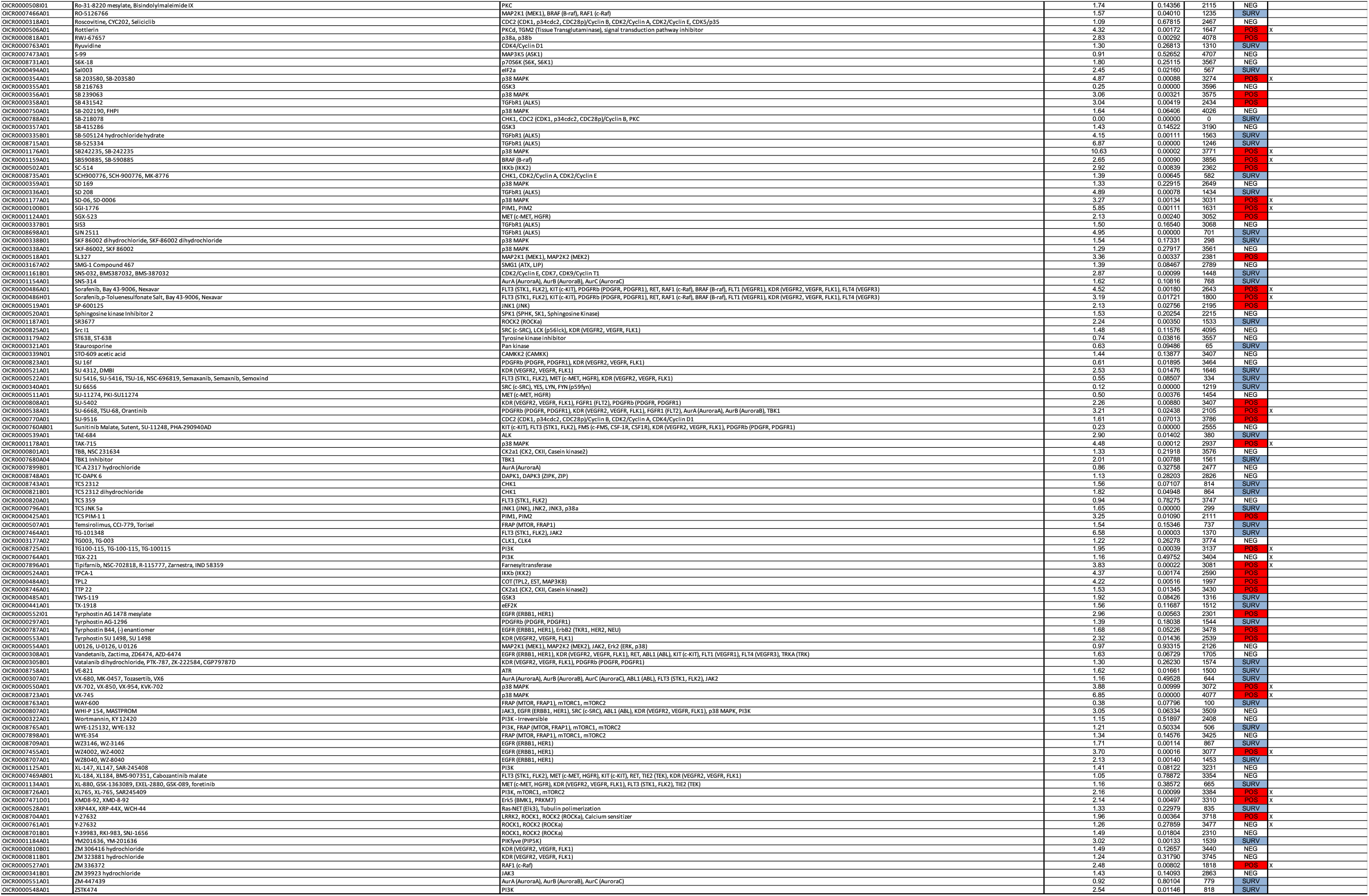

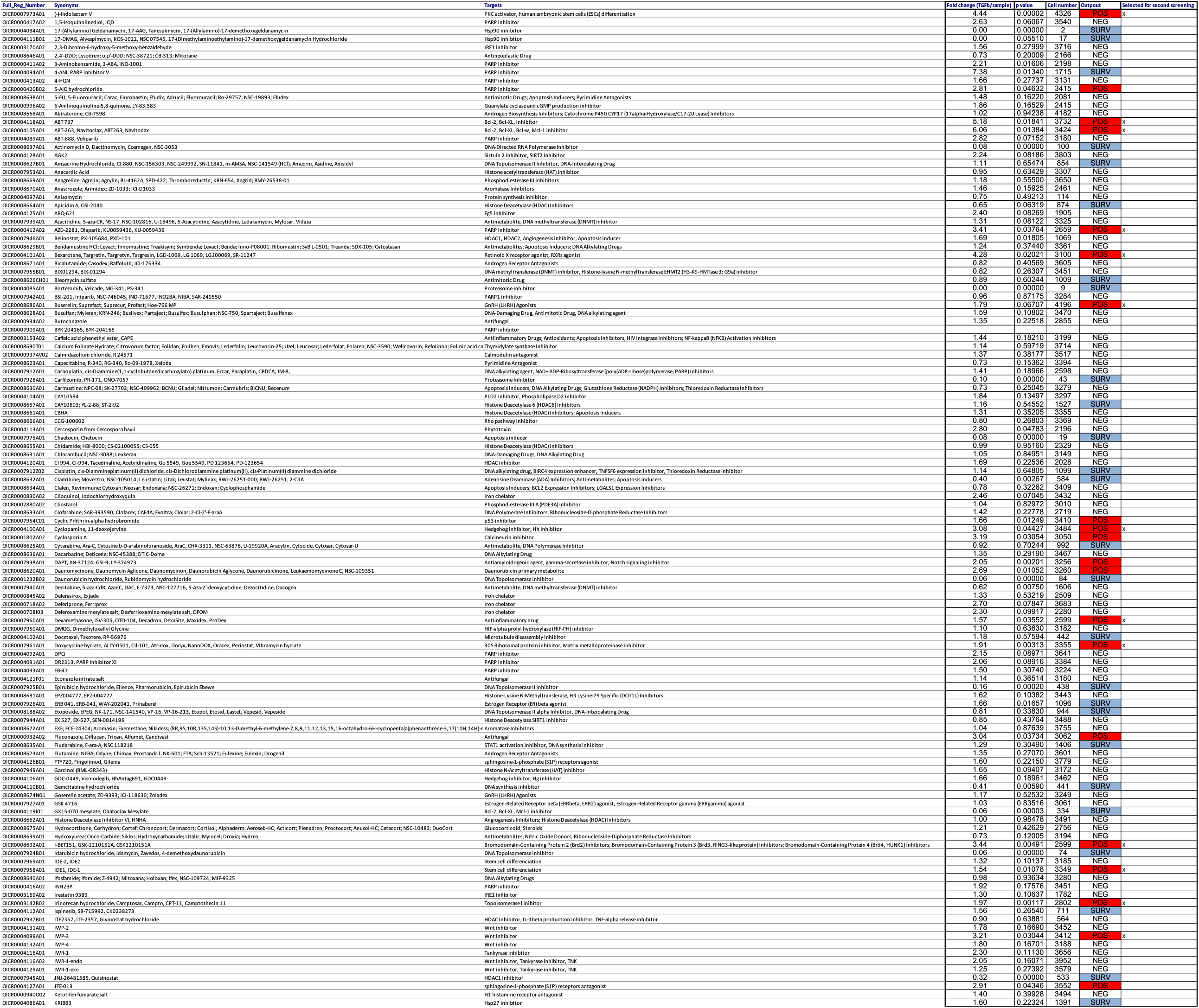
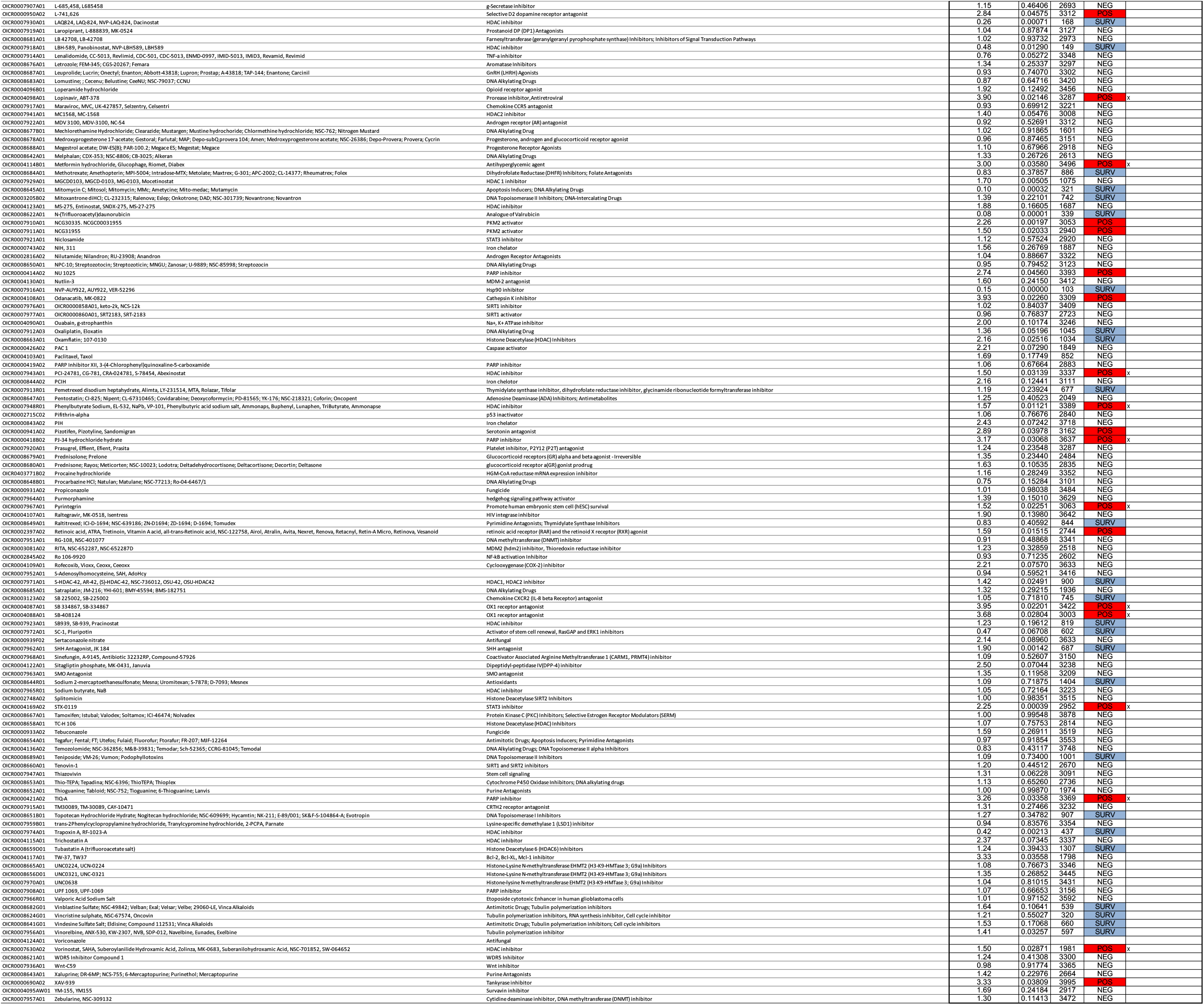

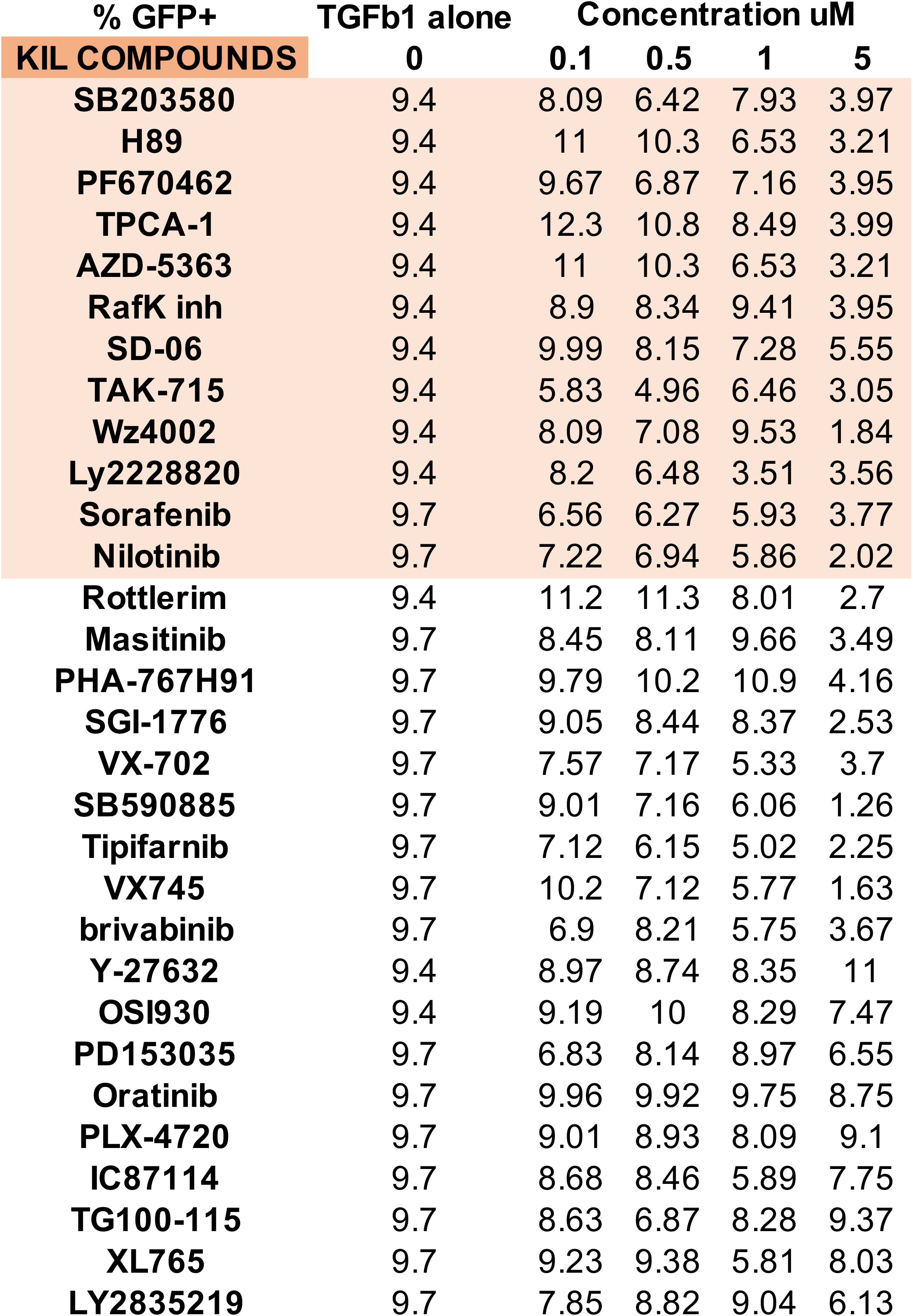

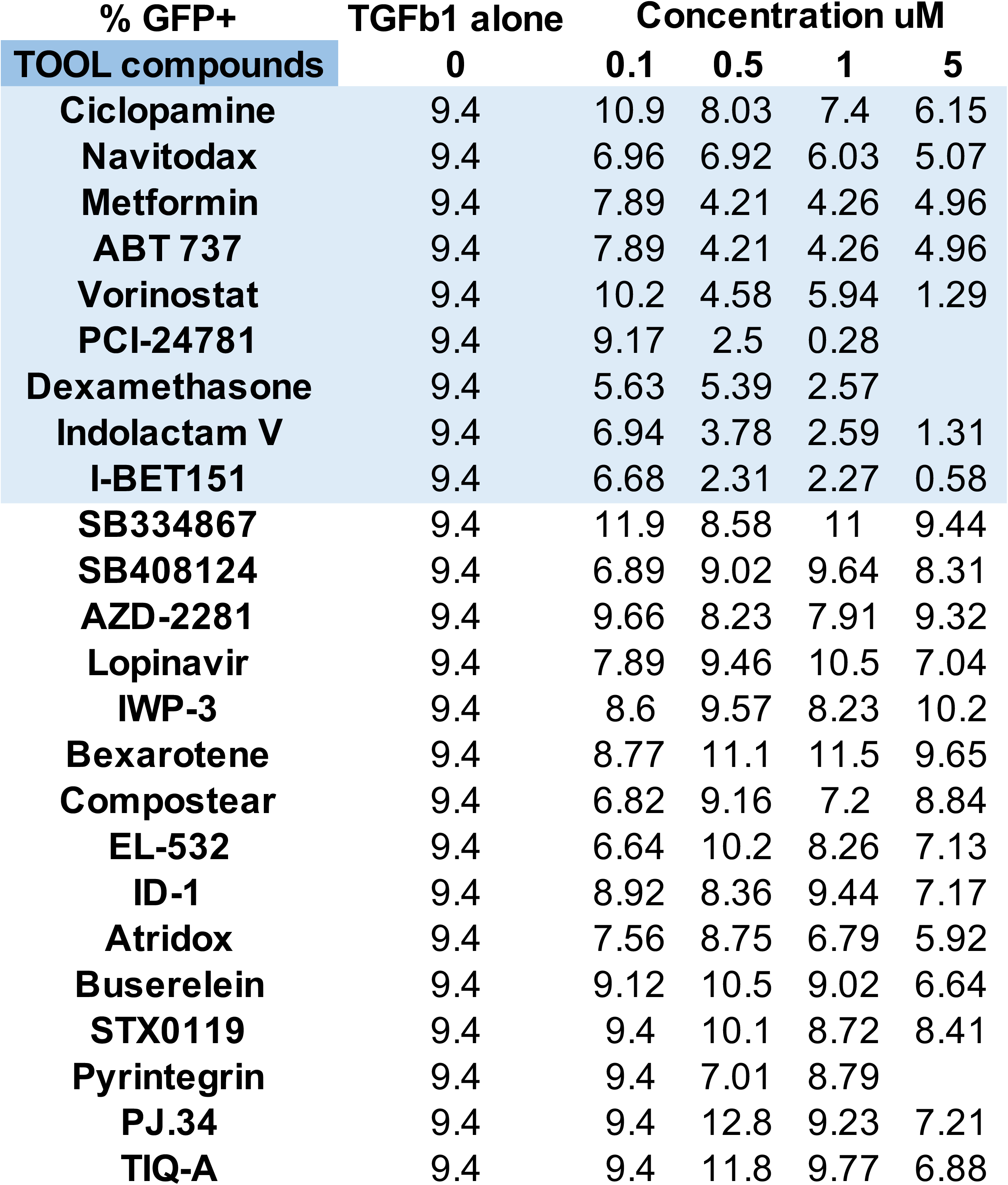

| Compound | Target | Fold change (sample/TGFb) | p value | Cell number | Output |
| --- | --- | --- | --- | --- | --- |
| A-366 | G9a/GLP | 1.04 | 0.80466 | 4511 | NEG |
| BAZ2-ICR | BAZ2A/B bromodomains | 0.78 | 0.16923 | 3555 | NEG |
| C646 | p300 | 0.80 | 0.31629 | 3848 | NEG |
| CL-994 | HDAC | 0.82 | 0.49564 | 2164 | NEG |
| GSK-LSD1 | LSD1 | 1.30 | 0.13926 | 3965 | NEG |
| GSK2801 | BAZ2A/B | 0.86 | 0.61931 | 3376 | NEG |
| GSK343 | EZH2 | 1.17 | 0.45595 | 2400 | NEG |
| GSKJ4 | JMJD3, UTX | 0.31 | 0.00128 | 895 | SURV |
| I-CBP112 | CBP/p300 | 0.58 | 0.06423 | 3461 | NEG |
| IOX2 | PHD2 | 0.90 | 0.54481 | 3816 | NEG |
| JQ1 | pan-BRD | 0.10 | 0.00032 | 2135 | POS |
| LAQ824 | HDAC | 0.45 | 0.89918 | 83 | SURV |
| LLY-507 | SMYD2 | 0.48 | 0.02312 | 2305 | POS |
| OF-1 | BRPR1-3 | 0.33 | 0.00780 | 318 | SURV |
| OICR-9429 | WDR5 | 1.01 | 0.95879 | 3953 | NEG |
| Olaparib | PARP1/2 | 0.69 | 0.05328 | 2871 | NEG |
| PFI-1 | BET | 0.29 | 0.00520 | 1677 | POS |
| PFI-2 | SETD7 | 0.56 | 0.01867 | 3544 | POS |
| PFI-3 | SMARCA | 1.38 | 0.03138 | 4468 | NEG |
| SGC-CBP30 | CREBBP/EP300 | 0.47 | 0.07700 | 3900 | NEG |
| SGC0946 | DOT1L | 0.90 | 0.57856 | 3512 | NEG |
| SGC707 | PRMT3 | 0.77 | 0.41633 | 3778 | NEG |
| UNC0638 | G9a/GLP | 0.88 | 0.46814 | 2892 | NEG |
| UNC0642 | G9a/GLP | 0.20 | 0.00105 | 3159 | POS |
| UNC1215 | L3MBTL3 | 0.81 | 0.27208 | 3855 | NEG |
| UNC1999 | EZH1/2 | 0.62 | 0.03198 | 2730 | POS |

## Notes

### Competing Interest Statement

The authors have declared no competing interest.

