## Supplementary figures and images for "Targeting fibrosis in the Duchenne Muscular Dystrophy mice model: an uphill battle"

### Figure S1

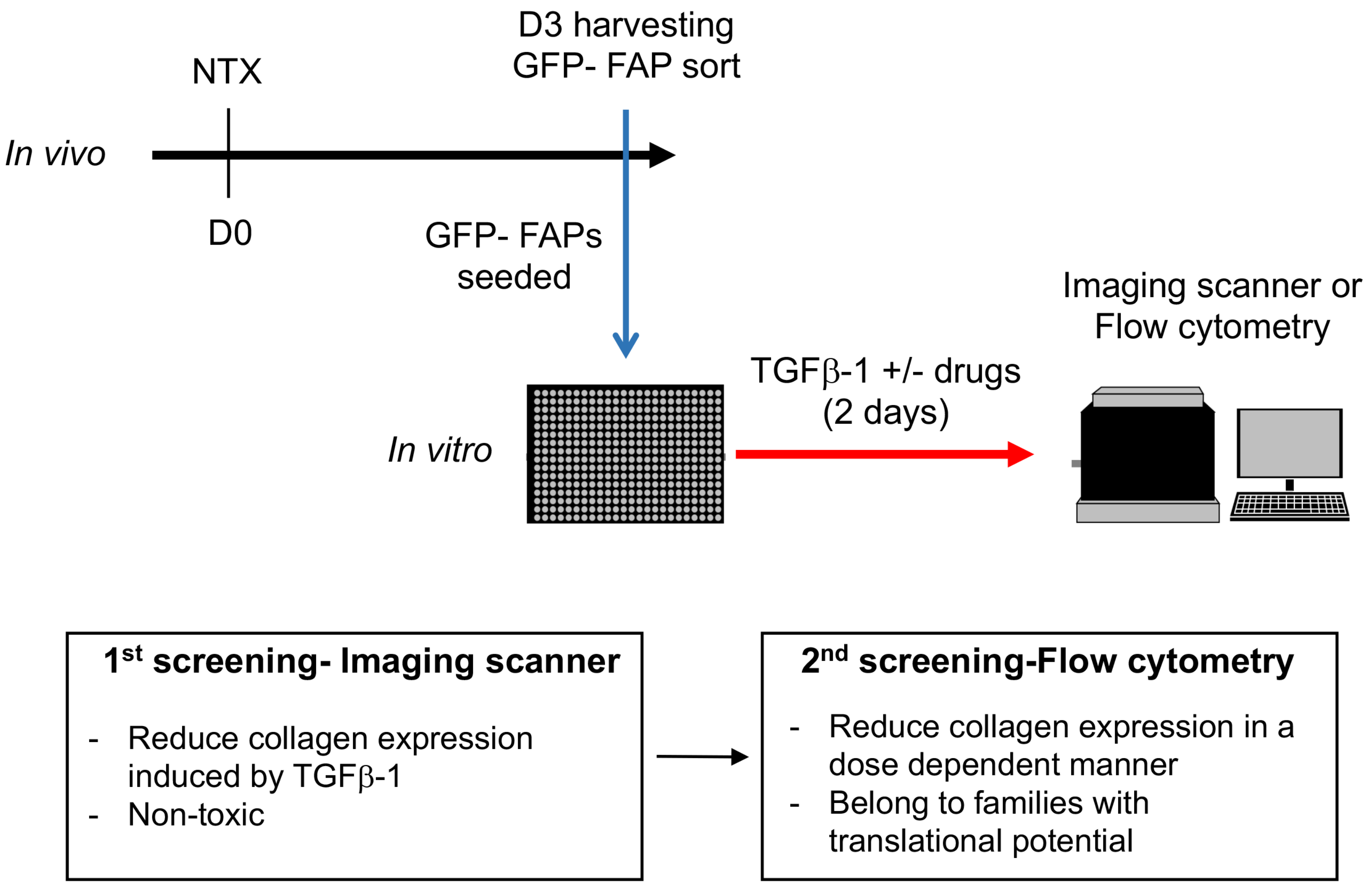

### Figure S2

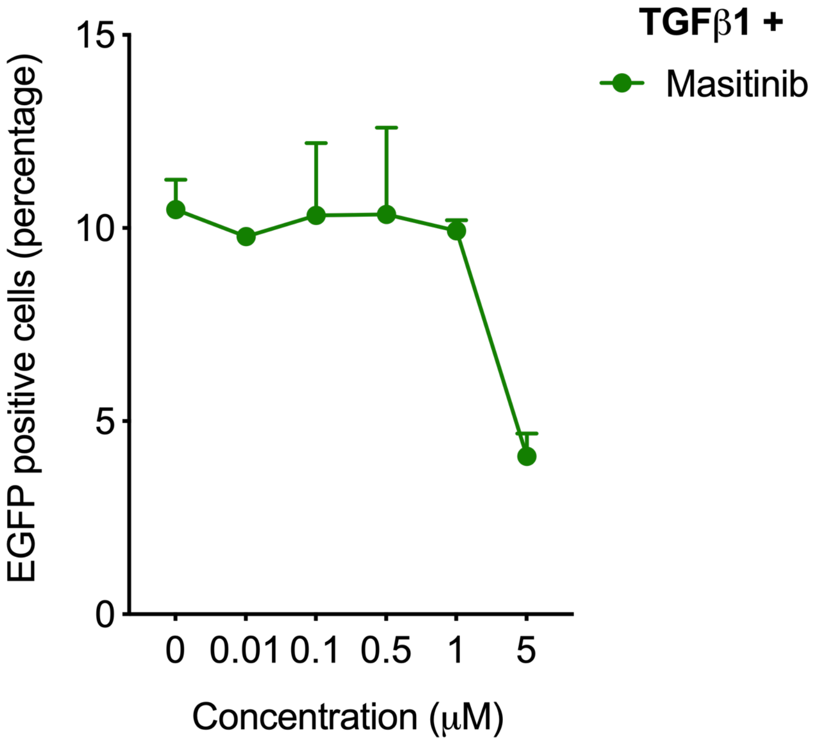

### Figure S3

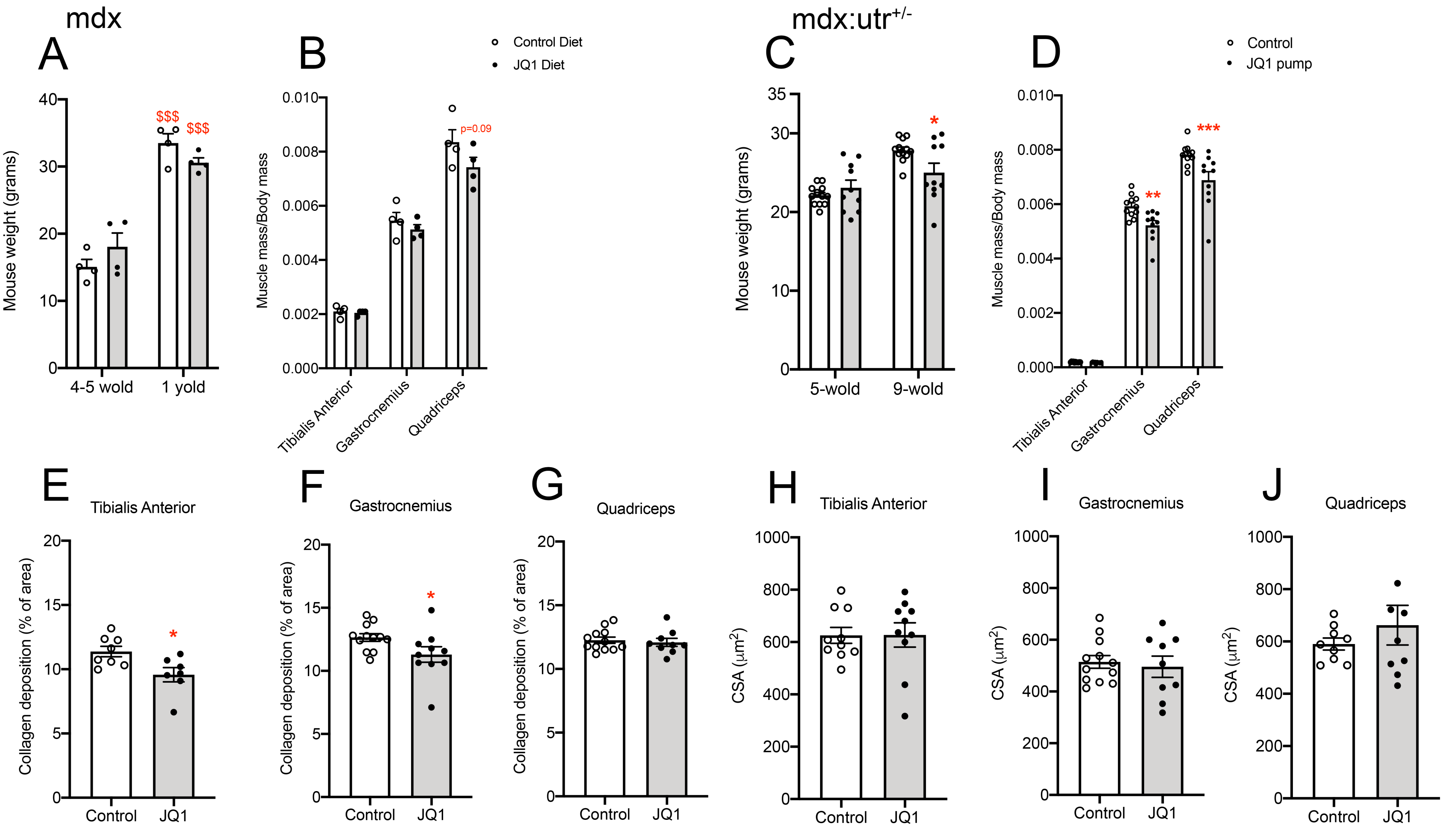

### Figure S4

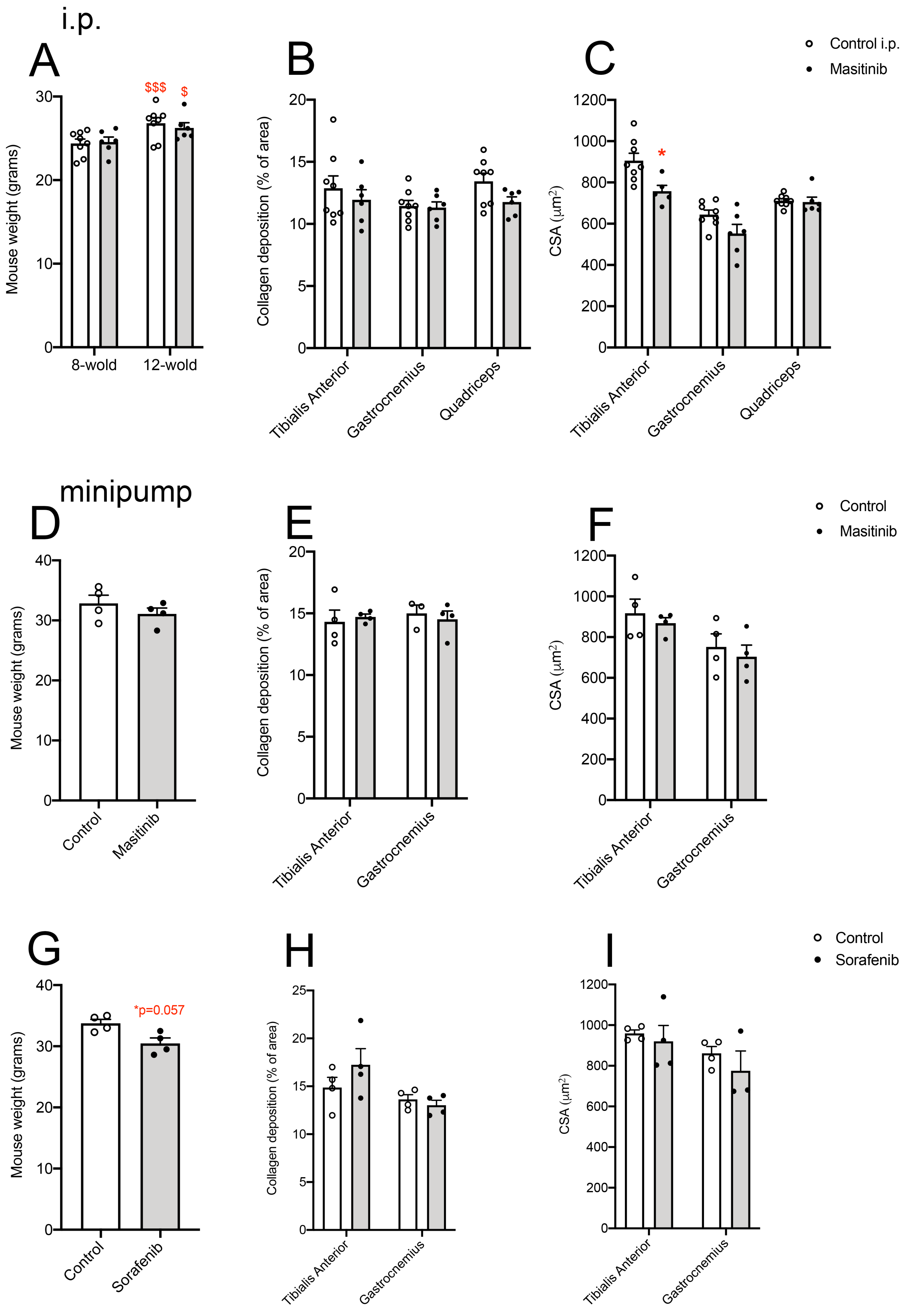
