## Supplementary material for "Targeting fibrosis in the Duchenne Muscular Dystrophy mice model: an uphill battle": Table S1

Table S1: Taqman probes

| Gene name | Reference |
| --- | --- |
| Collagen1a1 (*Col1a1*) | Mm00801666_g1 |
| Fibronectin1 (*Fn1*) | Mm01256744_m1 |
| Alpha-smooth muscle actin (*Acta2*) | Mm01546133_m1 |
| Periostin (*Postn*) | Mm01284919_m1 |
| Connective tissue growth factor (*Ctgf*) | Mm01192933_g1 |
| Hypoxanthine-guanine phosphoribosyltransferase (*Hprt*) | Mm00446968_m1 |
